# The role of IAA and its transport in the complex streptophyte algae *Chara braunii*

**DOI:** 10.1101/2024.10.16.618682

**Authors:** Katarina Kurtović, Stanislav Vosolsobě, Daniel Nedvěd, Karel Müller, Petre Dobrev, Vojtěch Schmidt, Piotr Piszczek, Andre Kuhn, Adrijana Smoljan, Tom J. Fisher, Dolf Weijers, Jiří Friml, John L. Bowman, Jan Petrášek

## Abstract

The role of auxin indole-3-acetic acid (IAA), a phytohormone with numerous morphogenic functions, is now well-established in land plants. Although the role of IAA and its transport in algae remains unclear, PIN-driven auxin export is probably an ancient and conserved trait within Streptophytes. Among streptophyte algae, *Chara* represents a genus with a considerable degree of complexity in body arrangement. This study investigates the auxin response and characterizes homologs of land plant PIN auxin efflux carriers in *Chara braunii*. Through regeneration experiments, we observed that IAA significantly promotes elongation of thallus and side branch development upon thallus tip decapitation, indicating an effect on morphogenesis. We show that IAA is actively uptaken and metabolized by thallus cells and that this process is influenced by N-1-naphthylphthalamic acid (NPA). To elucidate the underlying mechanisms, we cloned and sequenced the most expressed *Chara braunii* PINs, CbPINa and CbPINc. Using epitope-specific antibodies, we showed their presence in the plasma membrane (PM) of vegetative internodal cells and generative antheridial cells. Functional tests in tobacco BY-2 cells, supported by *in silico* docking of IAA and NPA to CbPINa and CbPINc, showed that PM-localized CbPINa interferes with auxin transport in contrast to ER-localized CbPINc. Finally, our phosphoproteome analysis indicated that IAA rapidly induces specific phosphorylation events, including RAF-like kinase phosphorylation, highlighting a potential role for IAA in fast signaling processes in *Chara braunii.* Altogether, we provide new insights into IAA role in *Chara braunii* morphogenesis and suggest that while the canonical auxin transport mechanism may not be conserved, auxin still may likely play a role in rapid signaling pathways in this close relative of land plants.

## Introduction

During the transition from water to land, plants underwent a series of developmental innovations leading to the establishment of a complex body (Harrison 2017; Donoghue et al. 2021; Bowman 2022). A key regulator of land plant development is the phytohormone auxin, which acts through local biosynthesis and directional transport resulting in formation of concentration gradients (Friml 2003). The directional flow of auxin, facilitated by the PIN family of auxin efflux carriers, is critical for plant morphogenesis and adaptive growth responses to environmental cues (Gao et al. 2008). Recent research has provided significant insights into the mechanisms of auxin action, including the structural elucidation of three PIN auxin efflux carriers (Ung et al. 2022; Su et al. 2022; Yang et al. 2022). However, the question of how and when auxin became a pivotal driver of morphological changes in land plants remains largely unanswered. This question cannot be fully addressed by studying land plants in isolation but requires a comprehensive investigation of their algal relatives (Skokan et al. 2019). Land plants (embryophytes) and streptophyte green algae, from which they emerged, together form the group Streptophyta (Becker and Marin 2009). Streptophyte algae comprise six clades, exhibiting significant morphological diversity within these clades (Buschmann 2020; Bierenbroodspot et al. 2024). Among these six clades, *Chara* spp. and *Nitella* spp., members of the Charophyceae family, possess the highest degree of complexity with respect to their body plan. Due to its large internodal cells, transparent gravitropic rhizoids, and rapid cytoplasmic streaming, *Chara* has been a model organism for decades, facilitating the study of fundamental cell biological processes (Kurtović et al. 2024).

Although genome sequencing revealed that *Chara braunii* lacks the auxin biosynthetic pathway involving the *TAA* and *YUCCA* genes, which convert Trp to a major auxin, indole-3-acetic acid (IAA) (Nishiyama et al. 2018), earlier and recent studies are consistently confirming the presence of IAA in biomass of various species of *Chara* and *Nitella* (Jahnke and Libbert 1964; Sztein et al. 2000; Hackenberg and Pandey 2014; Beilby et al. 2015; Schmidt et al. 2024), suggesting the possibility of an alternative biosynthesis pathway. In addition to the presence of auxin, its transport has also been studied in *Chara*. Prior to the availability of the *Chara* genome, Dibb-Fuller and Morris (1992) proposed that *Chara* might possess one or more functional efflux carriers. Their research, using radiolabeled IAA, demonstrated both auxin influx and efflux in *Chara* cells that were unaffected by the inhibitor N-1-naphthylphthalamic acid (NPA). Conversely, Boot et al. (2012) showed that NPA-sensitive polar auxin transport occurs in *Chara corallina*, facilitated by an efflux carrier, but they concluded that *Chara* probably lacks influx carriers. Later, PIN-like proteins were detected by immunolocalization using heterologous anti-AtPIN2 antibodies in the antheridial filaments of *Chara vulgaris* (Żabka et al. 2016). Finally, genome sequencing confirmed that *Chara* indeed possesses six homologs of PIN auxin efflux carriers, the highest number among all streptophyte algae (Hori et al. 2014; Cheng et al. 2019; Liang et al. 2020; Vosolsobě et al. 2020), at least five ATP-binding cassette B (ABCB) homologs, but no AUX/LAX influx carriers (Nishiyama et al. 2018).

Furthermore, exogenous IAA has been shown to promote rhizoid growth in decapitated *Chara* thalli of (Klämbt et al. 1992), affect ion transport (Zhang et al. 2016), induce transient depolymerization of microtubules (Jin et al. 2008), and to accelerate the process of differentiation of antheridial filament cells (Godlewski 1980). However, none of these studies further explored potential modes of IAA perception by *Chara*. The genome of *Chara* encodes certain elements of the auxin signaling pathway, in particular, an AUXIN RESPONSE FACTOR (ARF) and two Aux/IAA sequences, however, these components are not functionally equivalent to those in the canonical auxin signaling pathway of land plants (Nishiyama et al. 2018). The lack of canonical signaling pathway can be further supported by the absence of the TIR1/AFB1 receptor, which is an apomorphy unique to land plants (Bowman et al. 2017; Mutte et al. 2018; Su et al. 2023). On the other hand, *Chara* encodes a single ABP1 homolog (Carrillo-Carrasco et al. 2023), suggesting the possibility of a functional non-canonical pathway (Friml et al. 2022).

Given the ambiguous results in the literature and the availability of genomic information on *Chara braunii,* this study focuses on describing auxin responses and functional testing of *Chara* PIN homologs. We show that IAA treatments promoted side branching of regenerated thalli upon decapitation. To test the possible involvement of carrier-mediated transport in the *Chara* growth responses, we cloned two out of six PIN homologs, and showed that in *Nicotiana tabaccum* BY-2 cells CbPINa, but not CbPINc, influences the accumulation of radioactively labeled auxin. These results were supported by an *in silico* docking approach. Immunolocalization using specific antibodies showed that both CbPINa and CbPINc are associated with the PM in vegetative and generative cells of *Chara*. However, their expressions in *Arabidopsis thaliana*, and bryophyte *Marchantia polymorpha*, did not rescue the mutant phenotypes, despite their association with the PM and polar localization of CbPINa in gametangiophore stalks of Marchantia. To analyze rapid auxin action in *Chara*, we integrated phosphoproteomic analysis with cytoplasmic streaming assay, revealing IAA-specific changes in the phosphoproteome, including MAP4K homolog as a dominant target of IAA response and also the activation of a RAF-like kinase homolog. This provides evidence of conserved rapid auxin signaling in streptophytes, as reported by Kuhn et al. (2024). Additionally, our study demonstrates that IAA promotes ultrafast cytoplasmic streaming in germinating oospores, suggesting a link between auxin-mediated phosphorylation events and cytoplasmic streaming dynamics.

## Results

### IAA induces more axillary branches in nodes of decapitated *Chara* thalli

Although IAA effects on rhizoid growth promotion (Klämbt et al. 1992) and antheridia mitosis (Godlewski 1980) have been shown in previous studies, there has not been an attempt to identify the morphogenic role of IAA. Considering NPA-sensitive auxin transport has been previously shown in *Chara* (Boot et al. 2012), we tested whether external applications of IAA or NPA could cause morphological changes. We focused on the process of regeneration of new thallus upon decapitation, which is derived from nodal cells whose cell initials possess stem cell-like capacity (Heß et al. 2023). When decapitated explants containing two nodal complexes connected with one internodal cell were treated with 1 µM IAA, they regenerated longer thallus (Fig. 1a), while treatments with NPA had no effect. Furthermore, we tested if IAA is indeed inactive in restoring the apical dominance after the tip decapitation, as shown previously (Clabeaux 2011). After decapitation, we counted the number of axillary branches that developed from nodal cells at the base of thallus and found surprisingly that there were significantly more branches present after IAA application (Fig. 1b), and NPA again had no effect (Fig. 1b). As we have shown recently (Schmidt et al. 2024), there is evidence for biosynthesis and metabolism of IAA in *Chara braunii*. We were therefore interested in the fate of applied IAA and noted that it was completely depleted from the medium only 6h after treatment (Fig. 1c). In contrast, IAA significantly accumulated in *Chara* biomass, where it decreased during a 24h incubation to levels of DMSO-treated mock controls (Fig. 1c). We also observed the formation of IAA metabolites (Fig. S3). Interestingly, under NPA treatment IAA levels in the biomass after 24h almost depleted (Fig. S3), supporting indirectly the scenario where inhibited IAA efflux allowed its metabolization in cells. Altogether, these results suggest that exogenous IAA is imported by *Chara,* and it has morphogenic effects.

**Fig. 1.**
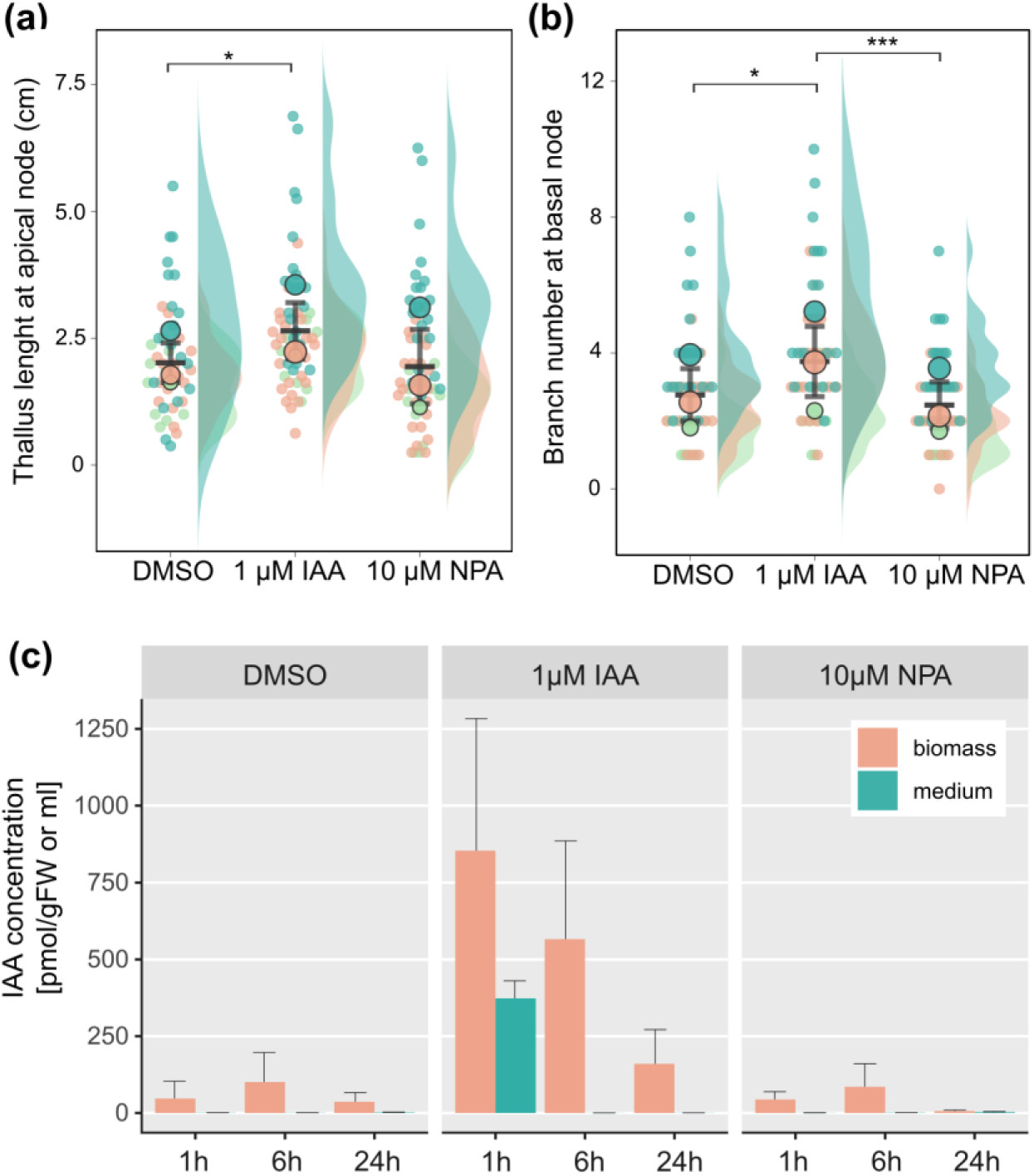
Decapitated *Chara* explants react to IAA and metabolize it quickly. **(a)** Newly regenerated thallus length upon treatments with DMSO mock control, 1 µM IAA, 10 µM NPA. *p* < 0.05 *. **(b)** Number of axillary branches at the basal node. *p* < 0.001 ***. Error bars in **a, b** show mean with SE, *n* = 10 (green), 20 (orange), and 20 (blue) Colors represent individual experiments. **(c)** Concentration of IAA in *Chara* biomass and media determined 1h, 6h, and 24h by LC/MS in mock (DMSO), 1 µM IAA and 10 µM NPA treated samples. Error bars indicate SE.

### Identification of six *CbPIN* paralogs and their expression

The effects of IAA on branching and NPA-specific changes in IAA levels suggested that specific auxin transporters could be involved in the auxin flow in *Chara.* Following the approach used in the functional analysis of PIN homolog from *Klebsormidium flaccidum* (Skokan et al. 2019), we aimed to clone PIN genes from *Chara braunii*. The whole genome sequencing revealed that *Chara braunii* has six PIN homologs: g29962, g29961, g41200, g50423, g50425 and g84230 (Nishiyama et al. 2018). We named the genes *CbPINa*, *CbPINb*, *CbPINc*, *CbPINd*, *CbPINe* and *CbPINf*, respectively (Table S1). Compared to land plant *PINs*, whose introns are a couple of hundred bases in length, *Chara braunii PIN*s are either intron-less (*PINa, PINb*, and *PINf*) or contain long introns (*PINc, PINd*, and *PINe*) from several thousand up to almost 20 thousand base pairs in length (Fig. 2a). *In silico* predictions of secondary protein structures of these two *Chara* PINs show that PINa has a long hydrophilic loop corresponding to the size the loop in canonical AtPIN1, while CbPINc has an extremely long cytosolic loop (>1000 amino acids) (Fig. 2b). Both *Chara* PINa and PINc have a canonical structure of ten highly conserved transmembrane domains (Fig. 2c). To analyze the expression patterns of individual *CbPINs*, we screened all *Chara braunii* publicly available RNA-seq data (Fig. S4b), which indicated preferential expression of *CbPINa* in protonema and rhizoids, while *CbPINc* was expressed in all analyzed stages besides rhizoids and oospore (Fig. S4b). We confirmed these results by RT-qPCR (Fig. S4a) from cDNA isolated from vegetative cells, where only *PINa* and *PINc* were successfully amplified and their identity verified by sequencing. Based on these results, we further focused on the functional characterization of *Chara PINa* and *PINc*.

**Fig. 2.**
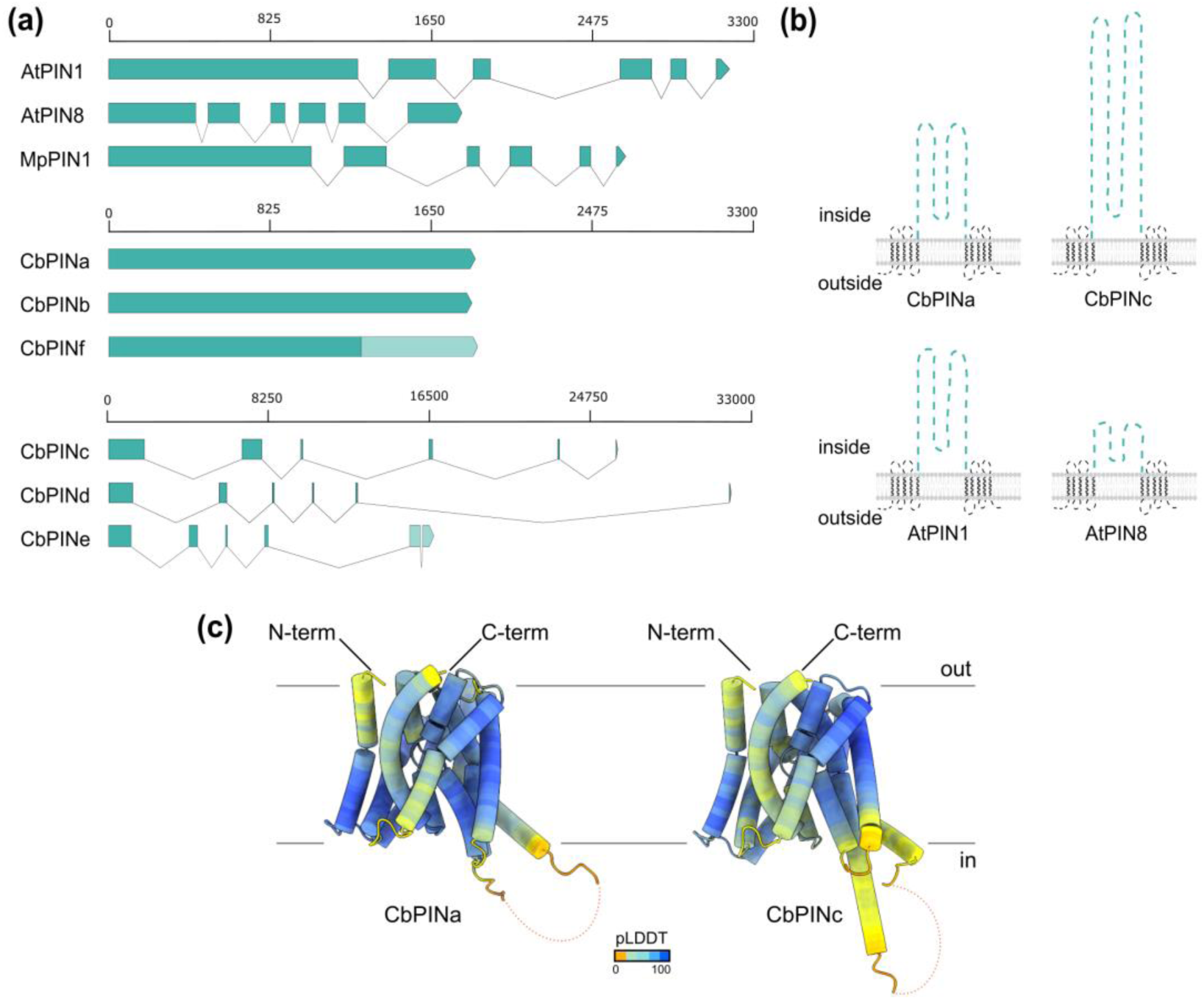
*Chara braunii* has six PIN homologs. **(a)** Exon-intron gene structures of *Arabidopsis thaliana PIN1 and PIN5*, *Marchantia polymorpha PIN1* and six *Chara braunii PIN*s, intron-less *CbPINa*, *b*, and *f*, and long intron-containing *CbPINc*, *d*, and *e*. Light green represents the missing regions or uncertain regions of the gene. **(b)** Schematic representation of secondary structures of CbPINa and CbPINc in comparison to the Arabidopsis AtPIN1 and AtPIN8. The predictions were performed using DeepTMHMM. **(c)** Predicted 3D structures of CbPINa and CbPINc proteins using AlphaFold. Color coding by pLDDT represents the confidence of the predicted structure compared to the true structure. Hydrophilic loop is unstructured and removed.

### *Chara braunii* PINs are localized in the charasome PM of internodal cells

NPA-sensitive auxin transport independent of cytoplasmic streaming has previously been demonstrated in internodal cells of *Chara corallina* (Boot et al. 2012). In land plants, this transport occurs via PM-localized PIN auxin efflux carriers (Ung et al. 2023). Therefore, we decided to localize *Chara* PINs in the internodal cells that form charasomes, convoluted PM domains (Chau et al. 1994). Since there are currently no reliable protocols for stable genetic transformation of *Chara* (Kurtović et al. 2024) we generated epitope-specific PIN polyclonal antibodies against CbPINa and CbPINc and confirmed their specificity by Western blotting (Fig. S5b). To identify charasome PM, we stained internodal cells with FM 1-43 (Schmölzer et al. 2011) and observed a typical clustered PM arrangement using confocal microscopy (Fig. 3b). Indirect immunofluorescence staining of both CbPINa and CbPINc followed by confocal microscopy showed specific dotted signals (Fig. 3c) that were reminiscent of charasomes. We did not observe any specific signals in the controls, including the omission of primary antibodies and or in samples probed with pre-immune serum (Figs. S5a, c). To further confirm the identity of the PM signal, we co-stained CbPINa and CbPINc with antibodies against H^+^-ATPase, which was shown previously to reside on charasomes (Schmölzer et al. 2011). The fluorescence signals of both CbPINa and CbPINc showed partial co-localization with H^+^-ATPase (Fig. 3d) in charasomes, indicating that two *Chara* PINs are indeed bound to specific regions of the PM. However, since the co-localizations were only partial (see zoomed-in insets in Fig. 3d), we suggest that *Chara* PINs may reside on other membrane compartments besides charasomes, either on non-invaginated PM or ER. Importantly, we did not observe any unspecific binding of antibodies to the cell wall (Fig. 3e). Overall, we show that *Chara* internodal cells PINa and PINc are preferentially localized within the charasome PM compartments, which may indirectly suggest their involvement in the auxin transport across the membrane.

**Fig. 3.**
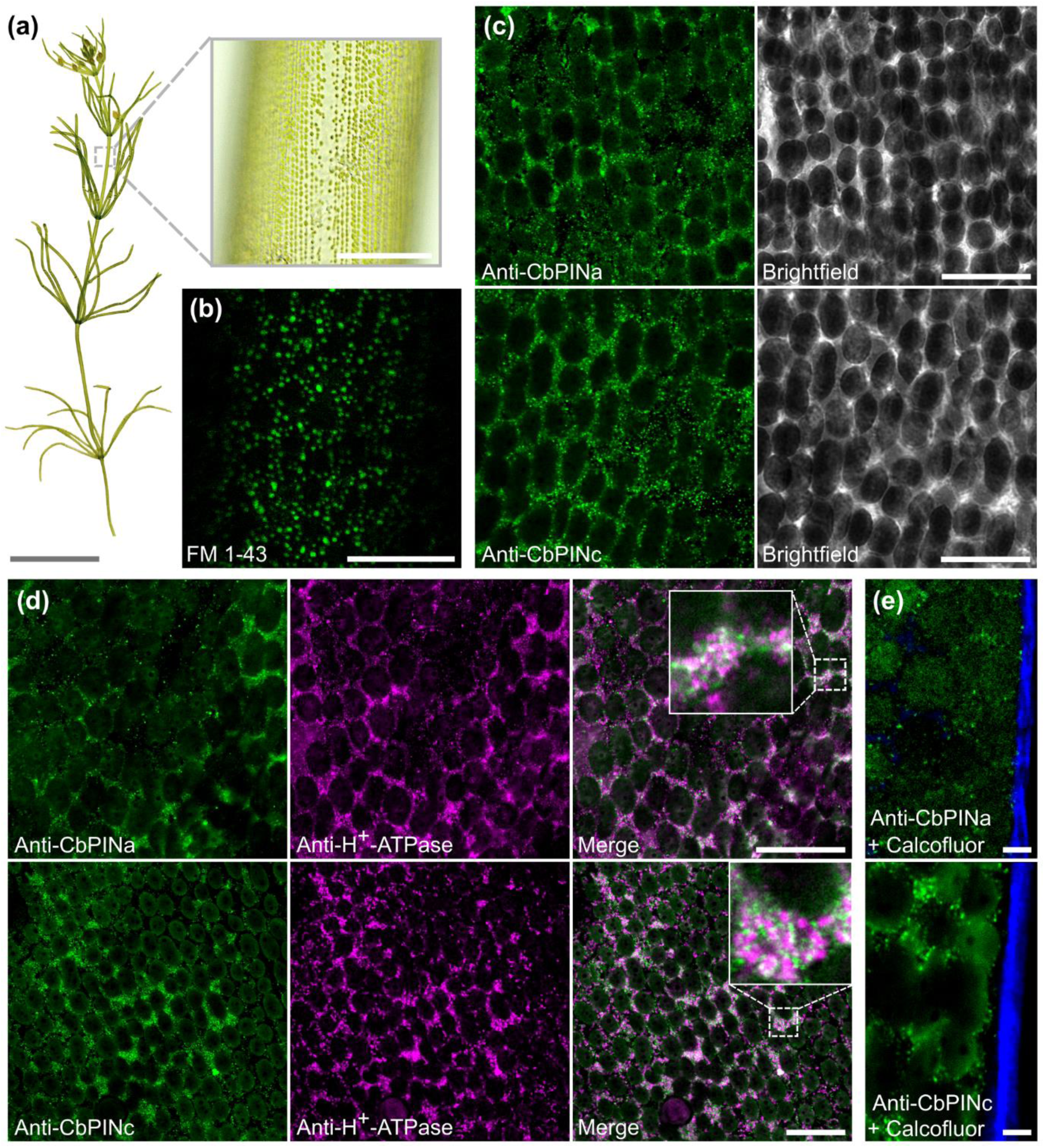
Immunolocalizations of CbPINa, CbPINc and H^+^-ATPase in the charasome PM of *Chara braunii* internodal cells. **(a)** Color brightfield image of *Chara braunii* thallus, strain S276, with zoomed-in internodal cell region **(b)** Confocal image of *in vivo* staining of internodal cell PM with FM 1-43. Green dots represent charasome PM while black dark ovals correspond to chloroplasts **(c)** Indirect immunofluorescence staining of CbPINa and CbPINc show the signal at the cell surface with its corresponding brightfield channel. **(d)** Co-localizations of CbPINa and CbPINc (green) with H^+^-ATPase (magenta) with charasome region zoomed-in insets. Note partial colocalization (white). **(e)** Immunofluorescence staining of CbPINa and CbPINc co-stained with calcofluor-white cell wall dye. Scale bars, 1 cm (thallus), 300 µm (zoomed-in inset) **(a)**, 20 µm **(b, c, d),** and 5 µm **(e)**.

### *Chara braunii* PINs are localized in the PM of antheridial cells

The analysis of *Chara braunii PIN* expression from publicly available RNA sequencing profiles showed that, besides thallus, *CbPINa* and *CbPINc* are also expressed in male generative organs, antheridia (Fig. S4). The PIN homolog presence in antheridia was also suggested by using heterologous antibodies against *Arabidopsis thaliana* PIN2 in *Chara vulgaris* (Żabka et al. 2016). The antheridia consist of generative antheridial filaments and non-generative shield cells, which encapsulate rosette-forming filaments (Fig. 4a). Antheridial filaments have a continuous PM that can be stained *in vivo* with FM 1-43 (Fig. 4b). Our immunolocalizations showed that CbPINa, but not CbPINc, is present in shield cells (Fig. 4c). In contrast, we localized both CbPINs at the edges of antheridial filaments (Fig. 4d), in a non-polar manner (zoomed-in insets in Fig. 4d). The signal was stronger towards the ends of filaments and was not present in capitular (progenitor cells), nor in manubria. We confirmed the identity of PM signal by co-staining with H^+^-ATPase antibodies, which showed that H^+^-ATPase does localize to the cell in the same regions as PINs at the edges of the cells, but the signal was also present inside cells, around the DAPI-stained nucleus (zoomed-in insets in Fig. 4e). We did not observe any specific signals in the controls, including the omission of primary antibodies and the inclusion of pre-immune serum (Fig. S5d), while immunostaining of α-tubulin served as a positive control (Figs. S6). Overall, the specific localization of CbPINa and CbPINc at the PM and their co-localization with H^+^-ATPase suggests that these proteins may play a role in the regulation of membrane-associated processes in male generative cells.

**Fig. 4.**
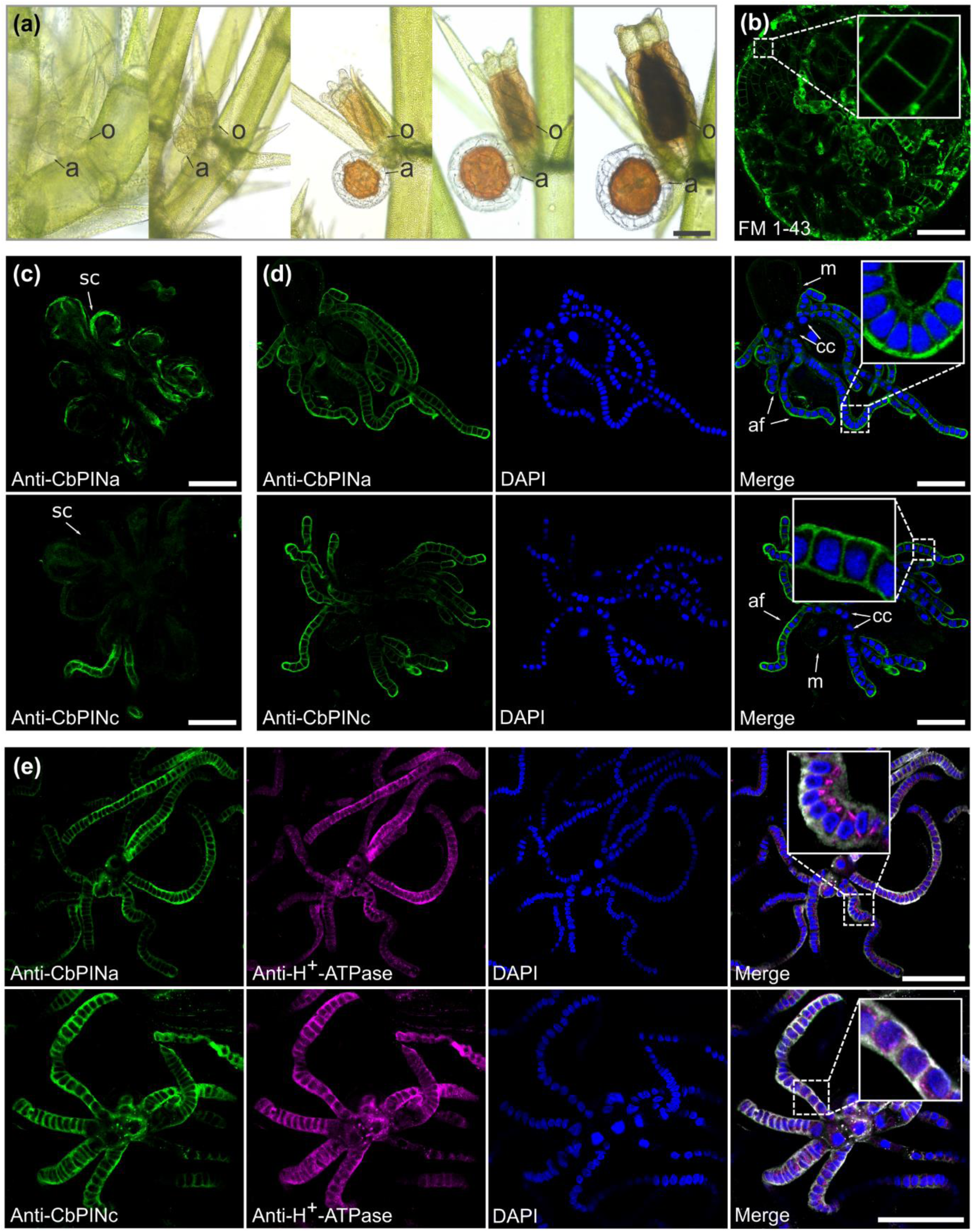
Immunolocalizations of CbPINa, CbPINc and H^+^-ATPase in the PM of antheridial cells of *Chara braunii*. **(a)** Progression of antheridium development on *Chara braunii* thallus. a, antheridium; o, oogonium. **(b)** *In vivo* staining of antheridium with FM 1-43. Zoomed-in inset shows the magnification of a filament with a smooth PM. **(c)** Differential presence of CbPINa and CbPINc in shield cells (sc), where CbPINa signal resides on the edges of the cell and there is no signal of CbPINc signal. **(d)** Antheridial filament stained with either CbPINa or CbPINc (green), DAPI staining of the nucleus (blue) and merged image with zoomed-in inset. m, manubrium, cc, capitular cells, af, antheridial filament. **(e)** Co-staining of CbPINa and CbPINc (green) with H^+^-ATPase (magenta) and nuclear staining with DAPI (blue). Merged images include all three channels and a zoomed-in inset of antheridial filament cells. Scale bars, 300 µm **(a)**, 100 µm **(b)**, 50 µm **(c-e).**

### CbPINa and CbPINc functional assays in tobacco BY-2 cells

To test the function of CbPINs, we introduced GFP-tagged *CbPINa* and *CbPINc* coding sequences under control of estrogen inducible XVE system (Zuo et al. 2000) and expressed them heterologously in tobacco BY-2 cells. Such XVE::CbPINa:GFP and XVE::CbPINc:GFP cell lines were used for confocal microscopy and auxin transport accumulation assays (Petrášek et al. 2006; Müller et al. 2019), following the setup that we used previously for testing PIN homolog from *Klebsormidium flaccidum* (Skokan et al. 2019). Upon induction, we observed a preferential localization of CbPINa at the PM, where it co-localized with FM 4-64 (Fig. 5a and S8a) and a weak signal at the ER. In contrast, CbPINc was predominantly localized to the ER, with a significantly lower PM localization compared to CbPINa (Fig. 5b). In the same induced cells, we performed auxin transport assays using radiolabeled 1-naphthaleneacetic acid ([^3^H]-NAA), a lipophilic synthetic auxin with very good specificity for auxin efflux carriers (Delbarre et al. 1996; Petrášek et al. 2006). Surprisingly, induced CbPINa showed higher accumulation of [^3^H]-NAA, almost reaching levels observed after the inhibition of endogenous auxin efflux with NPA (Fig. 5c and S8b). This suggests that CbPINa, while inactive as a carrier in tobacco cells, interferes with the activity of endogenous auxin efflux carriers, possibly by occupying their native residence in the PM and binding NAA. In contrast, induced CbPINc showed no difference in [^3^H]-NAA levels (Fig. 5c and S8b). Interestingly, despite CbPINa and CbPINc not showing auxin efflux activity in tobacco cells, multiple sequence alignment with land plant PINs (AtPIN1 and MpPIN1) (Fig. S7) revealed that they both possess a conserved phosphorylation site S2, which is crucial for regulating auxin efflux-dependent growth (Weller et al. 2017). To test the affinity of CbPINs to auxin and NPA, we performed an *in silico* docking experiment (Fig. 5d). The ligands, namely IAA, NAA and NPA interact with more amino acids when docked to CbPINa than with CbPINc. Furthermore, the predicted ligand binding affinities suggest that CbPINa has higher affinity to all three tested ligands compared to CbPINc (Table S2). In conclusion, our results suggest PM-localized CbPINa interferes with the PM auxin transport machinery, in contrast to ER resident CbPINc.

**Fig. 5.**
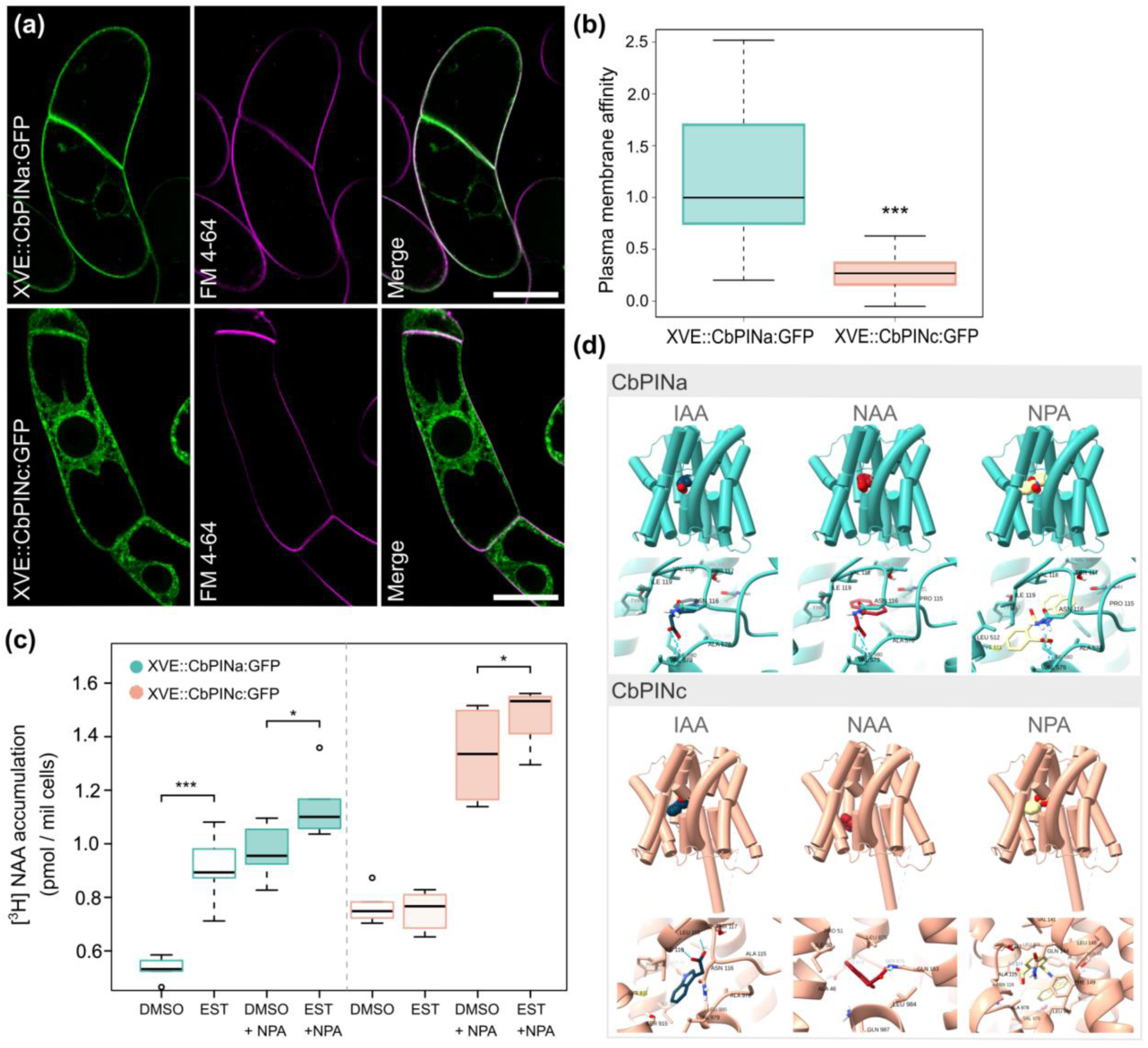
CbPINa localizes preferentially to the PM of BY-2 cells and interferes with [^3^H]-NAA accumulation while ER-localized CbPINc does not. **(a)** Induced XVE::CbPINa:GFP and XVE::CbPINc:GFP BY-2 cells. Shown are following; the GFP signal at the PM and ER in XVE::CbPINa:GFP and predominantly ER in XVE::CbPINc:GFP, FM 4-64 PM staining, and a merged image. **(b)** The comparison of *Chara* PINs PM affinity. The difference was calculated between 10 cells per variant, *p* < 0.001***. **(c)** [^3^H]-NAA accumulation in 5-day-old induced and non-induced XVE::CbPINa:GFP and XVE::CbPINc:GFP cells. NPA-sensitive NAA efflux was tested with the NPA (10 µM) applied at the beginning of the accumulation run. Data from two biological repetitions, with three technical repetitions. Each are shown with a median (black lines), p < 0.05*, p < 0.001*. **(d)** Ligand docking of IAA, NAA and NPA in *Chara braunii* PINa and PINc. 3D models show the position of the ligand within the protein and a close-up view shows the support sites with central residues highlighted. Predicted intermolecular hydrogen bonds are shown as dashed lines. Scale bar 25 µm **(a)**.

### *Chara* PIN heterologous expressions in Arabidopsis and Marchantia

Although the functional PIN auxin exporter from *Klebsormidium flaccidum* (Skokan et al. 2019) does not complement gravitropism defects when expressed under PIN2 promoter in Arabidopsis roots (Zhang et al. 2019), based on our results from tobacco cells, we decided to test the possible role of CbPINa and CbPINc in Arabidopsis. We first generated Arabidopsis *pin2* mutant plants expressing PIN2::CbPINa:GFP and PIN2::CbPINc:GFP (Fig. 6a). Using these lines, the non-polar localizations of both CbPINa and CbPINc in the PM and ER were observed in the root epidermal cells, while PIN2::PIN2:GFP showed a known shootward PM localization (Fig. 6a). These localizations corresponded to the results of gravitropic bending experiments, where CbPINa and CbPINc did not rescue the agravitropic phenotype of *pin2* mutant (Fig. 6b and c).

**Fig. 6.**
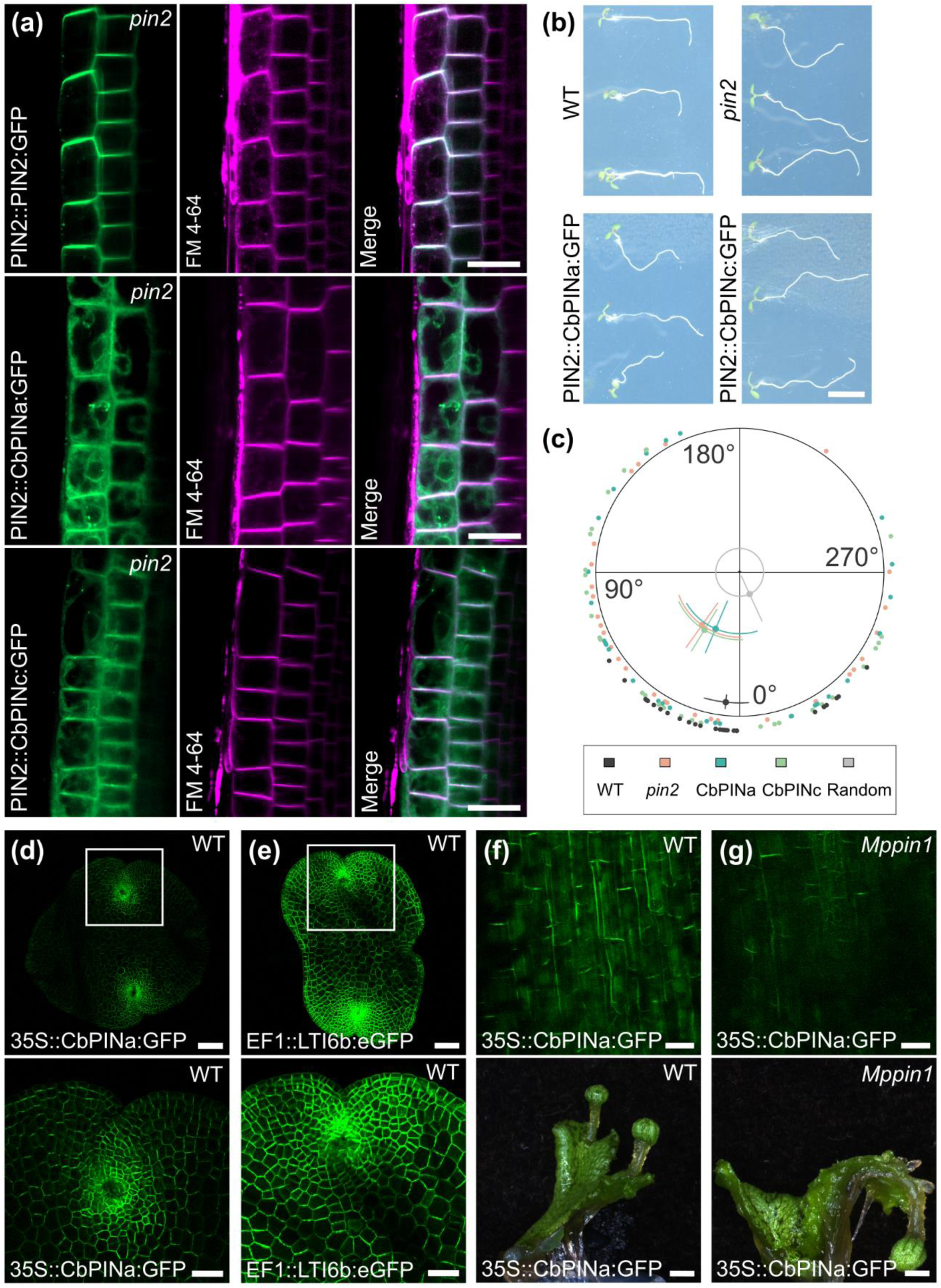
CbPINa and CbPINb do not complement Arabidopsis and Marchantia mutant phenotypes. **(a)** Localization of PIN2::PIN2:GFP/*pin2*, PIN2:CbPINa:GFP/*pin2* and PIN2:CbPINc:GFP/*pin2* in *Arabidopsis thaliana* root epidermal cells. Non-polar localization of CbPINa and CbPINa in the PM and ER, shootward AtPIN2 localization **(b)** Gravitropic root bending in 4-day-old seedlings of the WT, *pin2* mutant, and PIN2::CbPIN-expressing lines **(c)** Quantification of root bending assay. The diagram of the mean angle and length of the directional vector. The individual data points are shown on the outer area, while plotted data points are shown in the inner area. The confidence intervals for mean angle are indicated by an arc segment, while directional vectors are indicated by straight lines. **(d)** *Marchantia polymorpha* gemmaling (top) and zoomed-in gemmaling notch (bottom) expressing 35S::CbPINa:GFP/Wild-type. The GFP signal is predominantly present on plasma membrane. **(e)** EF1::LTI6b:GFP/Wild-type as a PM marker. Note the uniform localization across gemmaling. **(f)** Close up image of a female gametangiophore stalk expressing 35S:CbPINa:GFP/Wild-type (top) and white image of its corresponding of thalli with female gametangiophore (bottom). **(g)** Close up image of a female gametangiophore stalk expressing 35S:CbPINa:GFP/Mp*pin1* (top) and white light image of its corresponding thalli with female gametangiophore (bottom). Note the agravitropic stalk in the Mp*pin1* mutant background. Scale bars 20 µm **(a)**, 500 µm **(b)**, 100 µm (top) and 50 µm (bottom) **(d, e)** 50 (top) and 2 mm (bottom) **(f, g).**

In order to test the role of *Chara* PINs in a less complex organism, we expressed CbPINa::GFP under the control of 35S promoter in the bryophyte model *Marchantia polymorpha*, which contains just one canonical PM-localized PIN (Fisher et al. 2023). Interestingly, CbPINa localized predominantly to the PM in Marchantia gemmaling (Fig. 6d) and its distribution was, in comparison with the LTI6b PM marker (Fig. 6e), more concentrated on the gemmaling notches. Surprisingly, in female gametangiophore stalk CbPINa localized to the PM in a polar manner (Fig. 6f), however it did not complement the Mp*pin1* mutant phenotype (Fig. 6g). Altogether, although *Chara* PINs could be localized at PM in Arabidopsis and Marchantia, they do not complement the functions of canonical land plant PINs.

### IAA enhances cytoplasmic streaming and triggers global phosphorylation response in *Chara braunii*

Recent work has demonstrated that low concentrations of IAA trigger a specific and fast phosphoproteomic response in both streptophyte algae and land plants within two minutes of treatment. This response correlated specifically with the effect of low concentration of IAA on cytoplasmic streaming (Kuhn et al 2024). Therefore, we performed a set of observations of germinating oospores (Fig 7a) of *Chara* treated with increasing concentrations of IAA. The cytoplasmic streaming velocity increased in a concentration-dependent manner (between 1 and 100 μM IAA, and was stopped completely after 1 mM IAA (Fig. 7b, Videos S1-16). The streaming velocity was again restored after replacing the IAA solution with distilled water. Building on this observation, we conducted a phosphoproteomic analysis to understand if these fast IAA-triggered changes in cytoplasmic streaming are accompanied by similar IAA-specific phosphorylation modifications of proteins as they have been shown previously in *Klebsormidium* and *Penium* (Kuhn et al 2024). Furthermore, we used benzoic acid (BA) as control for the general effect of treatment by aromatic acid. Our results reveal that just two minutes treatment with 100 nm IAA induces specific shifts in both phosphorylation and dephosphorylation states of *Chara* proteome (Fig. 7c). While the number of detected sites was almost two thousand, the numbers of differential phosphosites were 114 in IAA vs. DMSO (Table S3 and S4), 116 in BA vs. DMSO and 107 in IAA vs BA, comparison at false discovery rate FDR ≥ 0.05. Furthermore, hyperphosphorylation upon IAA treatment represented the majority of differential phosphosites (55.10%), while upon benzoic acid, only 9.48% differential phosphosites were hyperphosphorylated. Furthermore, benzoic acid (BA) does not induce the IAA-like phosphorylation changes, as shown by a weak correlation between two treatments (Fig. 7d). Although *Chara* PINs were not detected in our dataset, we identified a distinct set of candidates potentially involved in auxin response. The most strongly influenced candidate is homolog of MAP4K class III (A0A388LCC7), dephosphorylated at Ser793 and Ser332. Notably, our data indicate that the *Chara* homolog of a RAF-like kinase (A0A388MAZ7) is phosphorylated by auxin at Ser1509, further confirming its deeply conserved evolutionary role in rapid auxin responses. To elucidate how the differentially phosphorylated candidates are involved in known and predicted protein-protein interactions, we performed a STRING analysis (Fig. S9), which revealed clusters of proteins involved in autophagy, transcriptional regulation, endocytosis, and cation transport, namely sodium/hydrogen exchanger (A0A388MCN6; hyperphosphorylated at Ser499).

**Fig. 7.**
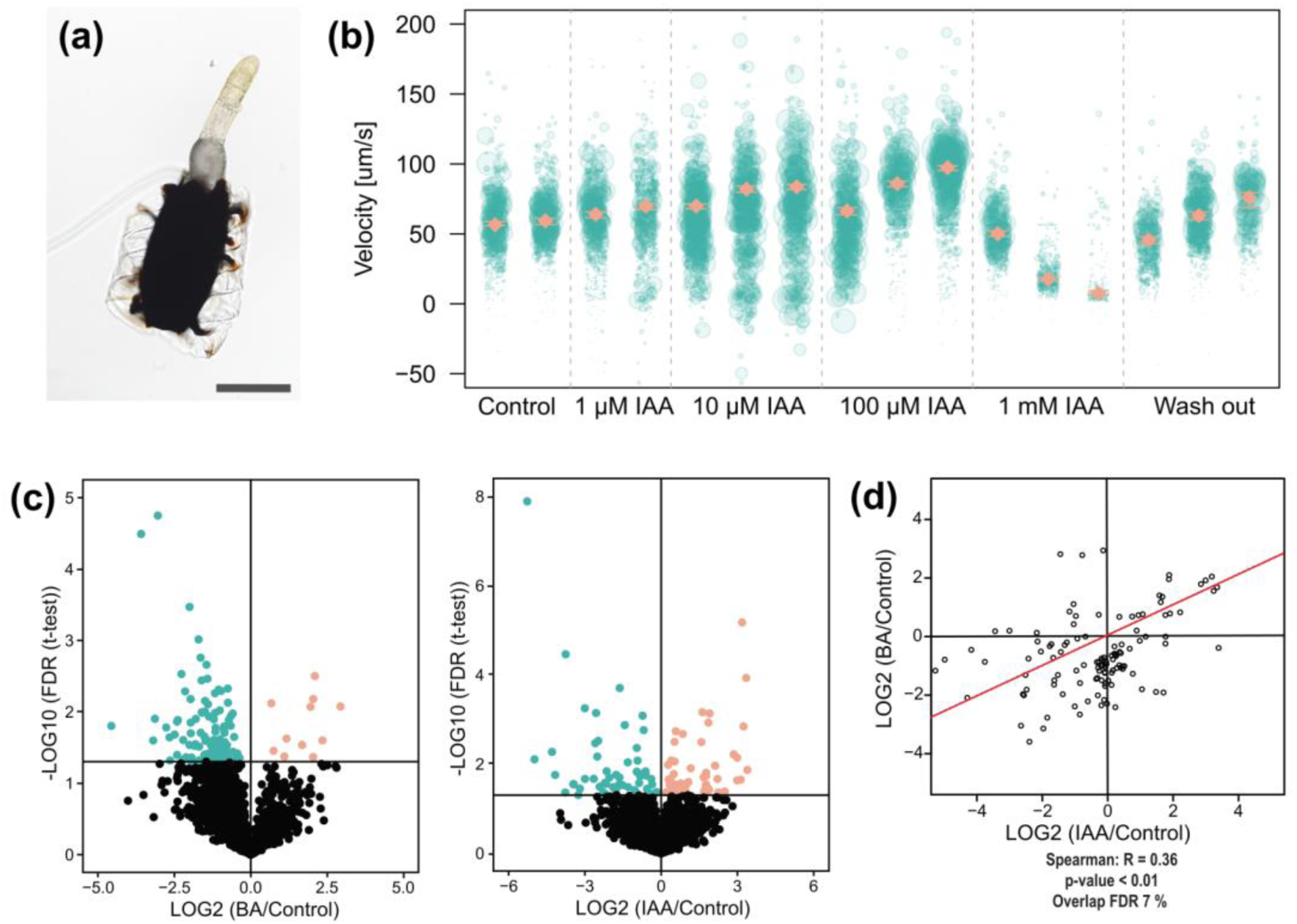
IAA enhances *Chara* cytoplasmic streaming and induces a specific phosphoproteomic response. **(a)** Brightfield image of a germinating oospore **(b)** Cytoplasmic streaming velocity in germinated oospore upon various concentrations of IAA treatment. Green points represent the speed of tracked particles. The orange square represents the weighted mean of velocity with 95% confidence intervals. **(c)** Distribution plots of significantly differential phosphosites comparing 2-minutes of 100 nM IAA treatment with DMSO treatment and 100 nM BA treatment with DMSO treatment. Orange color represents hyperphosphorylated phosphosites, blue hypophosphorylated, while black represents detected but not significant sites. **(d)** Plot comparing the differential phosphosites (FDR ≤ 0.05) after a 2-minute treatment with 100 nM IAA (x-axis) to the fold-change of the corresponding phosphosites following a 2-minute treatment with 100 nM benzoic acid. The red line represents the regression line. Scale bar 250 µm **(a)**.

## Discussion

### The response of *Chara braunii* to IAA

The study of the evolutionary origin of auxin biology and responses is one of the most fundamental tasks of plant evo-devo (Bowman et al. 2021). Auxin, primarilyIAA, is known to play a crucial role in regulating growth and development in land plants, particularly in processes such as apical dominance, cell elongation, and organogenesis (REF). Understanding how auxin responses originated and evolved sheds light on the evolution of plant complexity. In this context, our study explores the IAA role in *Chara braunii,* a late-diverging streptophyte alga, which has emerged as a model system for studying plant evolution (Holzhausen et al. 2022). Previous research showed that IAA promotes rhizoid growth (Klämbt et al. 1992) and branching induction after thallus tip decapitation in *Chara,* reflecting early forms of apical dominance (Clabeaux 2011). In this study, we tested the hypothesis that IAA influences the regeneration of new thalli following decapitation in *Chara braunii*. Our findings confirm that IAA-treated plants regenerated longer thalli, suggesting IAA role in growth promotion. However, unexpectedly, IAA induced promotion of axillary branches at the basal node, which contrasts with the typical auxin response in land plants (Fig 1b) where apical dominance suppresses lateral branching (Blake et al. 1983). Furthermore, the speed at which added IAA was depleted from the *Chara* medium and biomass suggests the presence of a highly effective auxin metabolic pathway (Fig 1c).

### *Chara* PINs are localized to the PM in vegetative and generative cells

To understand the mechanism behind the IAA response we examined the auxin genetic machinery present in *Chara*, focusing on characterization of *Chara* PIN auxin efflux carrier homologs. The PIN-FORMED (PIN) family of auxin efflux carriers is one of the most extensively studied membrane protein families in plants (Naramoto 2017), characterized by their ability to localize to different membrane compartments (Adamowski and Friml 2015). While endomembrane-localized non-canonical PINs are presumably involved in the regulation of intracellular auxin homeostasis (Sauer and Kleine-Vehn 2019), the canonical, plasma membrane-localized PINs play a crucial role in polar auxin transport, essential for plant growth and development (Wiśniewska et al. 2006). Moreover, the PIN homolog in the early-diverging Streptophyte *Klebsormidium flaccidum* has been shown as a functional auxin transporter (Skokan et al. 2019). The genome of *Chara braunii* (Nishiyama et al. 2018) revealed six PIN homologs (*CbPINa* to *CbPINf*), which is notable since more than two paralogs have not been observed in streptophyte algae (Vosolsobě et al. 2020). This likely resulted from specific gene radiation (Nishiyama et al. 2018), possibly driven by transposable elements, as suggested by the presence of unusually long introns (>20 kb) in its genome (Feng et al. 2024). We examined the structure and localization of two PIN homologs, CbPINa and CbPINc, which possess features similar to canonical PM-localized PINs in land plants (Bennett et al. 2014). Their cell-specific expression patterns (Fig. S4) suggest distinct roles, indicating potential involvement in *Chara* body plan development. As the absence of a transformation system in *Chara* limited our ability to directly test this hypothesis *in vivo*, we generated antibodies against CbPINa and CbPINc to study their localization. The localization patterns of both CbPINs in internodal cells, suggests a link to charasomes, plasma membrane structures involved in carbon uptake (Foissner et al. 2015), supported with a partial co-localization with H^+^-ATPase. In antheridia, CbPINa was present in shield cells, while both CbPINs were localized more abundantly at the plasma membrane towards the ends of filaments. This pattern contrasts with previous findings using heterologous antibodies against AtPIN2 in *Chara vulgaris*, which showed more signal in capitular cells (Żabka et al. 2016). Additionally, we found no signs of polar distribution of CbPINs in antheridial filament cells, further emphasizing the functional divergence between *Chara* PINs and their land plant counterparts.

### CbPINs are not functioning as auxin efflux carriers in heterologous systems

To assess the auxin transport capabilities of CbPINa and CbPINc, we expressed these proteins in tobacco BY-2 cells under an estradiol-inducible promoter. Unlike the PIN homolog in *Klebsormidium,* where induced KfPIN showed clear export in BY-2 cells (Skokan et al. 2019), we observed more ambiguous results with *Chara* PINs. Specifically, induced plasma membrane-localized, CbPINa showed higher accumulation of [^3^H]-NAA than non-induced, while CbPINc, primarily localized to the endoplasmic reticulum, did not significantly affect auxin transport. The lack of transport capacity may be due to several reasons. *Chara* PINs may have evolved to transport different form of auxin than tested in our transport assays. Furthermore CbPINs may require specific interacting proteins that are absent in BY-2 cells or post-translational modifications (e.g. phosphorylation), thus impairing their activity. Additionally, the extended hydrophilic loop of CbPINc (Fig. 2b) may lead to improper folding when expressed heterologously, potentially resulting in an autoinhibitory effect. We further extended our investigation to land plant models *Arabidopsis thaliana* and *Marchantia polymorpha*. Both CbPINs showed signal in the endoplasmic reticulum and plasma membrane, but unlike the polar AtPIN2, they did not localize polarly and failed to rescue the *pin2* mutant, indicating functional divergence from land plant PINs. In *Marchantia polymorpha* gemmalings, CbPINa predominantly localized to the PM, as confirmed by a PM marker, but failed to rescue Mp*pin1* mutant phenotype. These findings suggest CbPINa localizes the PM in both heterologous and homologous systems, while CbPINc is primarily ER-localized in heterologous systems but PM-localized in *Chara*.

### Rapid responses to IAA and its perception in *Chara*

While the nuclear auxin pathway is widely regarded as a feature unique to land plants that emerged during the process of terrestrialization (Hernández-García et al. 2024), recent discoveries of a conserved rapid auxin response across both plants and streptophyte algae suggest the existence of a more ancient auxin signaling pathway (Kuhn et al. 2024). Our analysis of *Chara* phosphoproteome coupled with the analysis of cytoplasmic streaming in germinating oospores revealed a distinct phosphorylation patterns. This includes identifying candidates like the MAP4K homolog of Arabidopsis BLUS1/TOT3/TOI (Pan et al. 2021), a sodium/hydrogen exchanger homologous to NHX5 and NHX6, which are essential for PIN6-mediated auxin homeostasis in Arabidopsis (Lv et al. 2020), and a RAF-like kinase homolog (Kuhn et al. 2024), all of which support the deeply conserved role of auxin in rapid cellular responses and are consistent with findings in other species. However, the number of significantly hyper- and hypophosphorylated phosphosites was smaller than in previously reported species (Kuhn et al. 2024), likely due to the limited coverage of the *Chara* genome or due to heterogeneity of samples.

Our results taken together suggest that while CbPINa and CbPINc may not act as conventional auxin transporters in heterologous systems, they may play a role in mediating auxin within *Chara braunii*. The presence of six PIN homologs hints at potential redundancy (Vieten et al. 2005) or specialization, warranting further study. Additionally, our findings reveal both conserved and novel auxin effects in *Chara*. The divergence in auxin responses, such as the promotion of axillary branching, suggests that auxin functions may have undergone significant evolutionary shifts. Future research should investigate the remaining four PIN homologs and the fast auxin response pathways to understand their role in plant complexity evolution.

## Materials and methods

### Plant materials and growth conditions

Two strains of *C. braunii* were used in this study: S276 from MAdLand (https://madland.science/.) and NIES 1604 from the NIES collection (https://mcc.nies.go.jp/). Both strains were grown on modified double-autoclaved soil-sand medium (SWCN-4) (https://mcc.nies.go.jp/medium/en/swcn4.pdf), supplemented with sterile distilled water, and propagated biweekly. The strain NIES 1604 was used for regeneration, germination and metabolic profiling, while strain S276 was used for DNA and RNA isolation and phosphoproteomic analysis. Cultures were maintained under fluorescent light (F15W/T8 AQUASTAR, cat num. 0002224; Sylvania) at room temperature, while regeneration experiments were conducted in a custom box with LED strips (construction of the box is detailed in methods S1 and Fig. S1), at 19-22°C, and a 14h light /10h dark cycle. To germinate oospores, mature oospores were harvested, air-dried for one month and stratified at 4°C for at least 4 months to break the dormancy. Before germination, oospores were sterilized with 1% Tween 20, followed by 8 min treatment with 10% commercial bleach, and rinsed thoroughly. Oospores were sown on 0.5% plant agar or in glass tubes with *Chara* medium (described above). The tobacco line BY-2 (*N. tabacum* L. cv Bright-Yellow 2) cells were cultured in a liquid medium containing 3% (wt/vol) sucrose, 4.3 g L^−1^ Murashige and Skoog (further referred to as MS) salts (Sigma-Aldrich, M5524), 100 mg L^−1^ inositol, 1 mg L^−1^ thiamine, 0.2 mg L^−1^ 2,4-dichlorophenoxyacetic acid, and 200 mg L^1^ KH_2_PO_4_ (pH 5.8). Cell suspensions were maintained on an orbital shaker (100 rpm) in darkness at 26°C and subcultured weekly. BY-2 calli stock was grown on the same media solidified with 0.6% (wt/vol) plant agar and subcultured biweekly. *A. thaliana* wild type Columbia-0 (Col-0) and *pin2* mutant were grown at 22°C with 16 h light/8h dark cycle on ½MS medium (Murashige and Skoog 1962). Marchantia gemmae were grown on 1.4% agar (A111, Phytotech Labs) containing ½B5, at 21°C under constant white light (45 µmol m^−2^ s^−1^ over the waveband 400-750 nm). Marchantia gametangiophore stalks were induced by supplementing with continuous far-red (FR) light (Quantum Devices Inc., Barneveld, WI, USA; 20–40 µmol m^−2^ s^−1^ over the waveband 715–745 nm).

### Chemical treatments

For regeneration experiments, decapitated thallus segments were treated with 1 µM IAA (indole-3-acetic acid), 10 µM NPA (N-1-naphthylphthalamic acid), and the equivalent amount of solvent DMSO (dimethyl sulfoxide), immediately upon propagation. Treatments were repeatedly added to *Chara* medium every 48h, until the harvest. For metabolic profiling, the treatments were performed on 3-weeks old *C. braunii* cultures. Individual plants were treated with 1 µM IAA, 10 µM NPA, and the equivalent amount of solvent (DMSO). The samples were collected 1h, 6h, and 24h after treatments in four replicates. For cytoplasmic streaming analysis, the agar-germinating oospores were carefully excised from the medium, placed on microscopic glass with drop of distilled water. The oospores were then treated with IAA ranging from 1 μM to 1000 μM IAA or with distilled water. The analysis of phosphoproteome upon treatments was peformed on 3-week-old cultures treated with 100 nM IAA, 100 nM benzoic acid (BA), or an equivalent volume of solvent (DMSO) as a negative control for 2 minutes.

### Nucleic acid isolation and RT-qPCR

About 50 mg of *C. braunii* thallus, strain S276, was harvested for DNA extraction. The DNA was extracted using the QIAGEN DNeasy Plant Mini Kit and concentration was measured with NanoDrop (Thermo Fisher). RNA was isolated using the FavorPrep Plant Total RNA Mini kit (Favorgen Biotech Corp) according to the manufacturer’s guidelines. DNase treatment was carried out using the DNase set (Macherey-Nagel). For RT-qPCR, approximately 1 µg of DNase-treated RNA was reverse-transcribed using M-MLV reverse transcriptase, RNase H-, point mutant (Promega). Quantitative real-time PCR was performed using GoTaq qPCR Master Mix (Promega) at 58°C annealing temperature on a LightCycler480 instrument (Roche). PCR efficiency was estimated using serial dilution of template cDNA. *Chara* homolog of elongation factor 1a (Genbank Acc. Nr. AB607260.1) was used as a reference gene. Positive transcript levels and the quality of PCR products were verified by melting curve analysis.

### Vector construction

The *Chara braunii* PINa coding sequence was amplified via PCR from genomic DNA (primers listed in Source Data). Since the original sequence was truncated at the N-terminus, a forward primer was designed 327 nucleotides upstream. The updated sequence was submitted to GenBank (PQ421084). For CbPINc, the first exon was amplified, and the rest was synthesized (Biocat, Germany). The accession number is in Table S1. Partial sequences were combined by PCR to create a full CbPINc sequence. GFP gene (no stop codon) was inserted at the 429^th^ amino acid for CbPINa and 558^th^ for CbPINc. Constructs were cloned into pJET1.2, transformed into *E. coli* (strain JM109), and subcloned into beta-estradiol-inducible vector pER8-XVE. To generate plasmids for genetic complementation in Arabidopsis, CbPINa:GFP, CbPINc:GFP fusions, and 1.4 kb PIN2 promoter were separately cloned into the Gateway entry vector pDONR221 and pPONRP4P1r vector. Constructs were later fused and cloned into Gateway destination vector pB7m24GW.3 by LR reaction. For genetic complementation in Marchantia, the Cauliflower Mosaic Virus (CaMV) promoter pro35S was cloned into HindIII/XbaI-cut pMpGWB401 (Ishizaki et al. 2015) as described in (Fisher et al. 2023). This destination vector (pro35S:GW GWB401) was recombined with CbPINa:GFP pDonor221 using LR clonase II (Invitrogen) to produce 35S:CbPINa:GFP GWB401. Lti6b:eGFP was amplified from a level 1 plasmid containing Lti6b:eGFP as a DNA template (Pollak et al. 2019). The fragment was subcloned into EcoRI-cut pENTR2b plasmid using a NEBuilderHiFi Gibson Assembly Kit (New England Biolabs, Ipswich, MA, USA). The Gateway destination vector pMpGWB403 (Ishizaki et al. 2015) was recombined with pENTR2b using LR clonase II (Invitrogen) to produce proEF1::Lti6b:eGFP GWB403.

### Genetic transformation

BY-2 cells were transformed by co-cultivation, according to the basic transformation protocol of (An 1985). Vectors carrying XVE::CbPINa:GFP or XVE::CbPINc:GFP were introduced into wild-type BY-2 and cultured in MS medium with 100 μg/mL cefotaxime and hygromycin for selection. (Petrášek et al. 2003). Transgene expression for microscopy and accumulation assays was induced by adding 2 μM estradiol during inoculation. Transgenic Arabidopsis plants were generated using the floral dip method and selected on solid, ½ MS medium containing 15 mg/mL of Basta (Glufosinate). Sporophytes of *Marchantia polymorpha* (ssp*. ruderalis*) (Causse 1993) were collected from a laboratory cross between Mel-1 and Mel-2 plants, male and female wild-type lines from a Melbourne population, Australia (Flores-Sandoval et al. 2015). For Mp*pin1* sporeling transformations, sporophytes were collected from a laboratory cross between Mp*pin1-4* and Mp*pin1-2* plants, as described in (Fisher et al. 2023). Transformation of *M. polymorpha* sporelings followed the description by (Ishizaki et al. 2008) using sporeling liquid media ½B5 (G398, Phytotech Labs) (Gamborg et al. 1968).

### Western blot and immunohistochemistry

Protein extraction, Western blot, and of immunolocalization of *Chara* PINs and H+-ATPase in internodal cells were performed using modified protocol from Schmölzer et al. (2011). The antheridia were immunostained according to (Żabka et al. 2016). Microtubules were stabilized with taxol and stained according to (Wasteneys et al. 1989). Detailed protocols are described in Supplementary Methods S2-S6.

### Protein extraction, phosphopeptide enrichment, mass spectrometry and data analysis for phosphoproteome

Detailed protocols are described in Supplementary Methods S7-S11.

### Determination of IAA levels and metabolites using LC/MS

Between 10-30 mg of treated *Chara braunii* thalli (treatment described above) was harvested and immediately frozen in liquid nitrogen. The samples were prepared according to Schmidt et al. (2024). Hormone analysis was performed with an LC/MS system consisting of UHPLC 1290 Infinity II (Agilent, Santa Clara, CA, USA) coupled to 6495 Triple Quadrupole Mass Spectrometer (Agilent, Santa Clara, CA, USA).

### Radio-labelled auxin accumulation assay

For accumulation assays, 5-day-old estradiol-induced and non-induced XVE::CbPINa:GFP or XVE::CbPINc:GFP BY-2 cell suspensions were prepared by filtering the liquid phase and resuspending the cells in uptake buffer (Petrášek et al. 2003). Cells were incubated in the dark for 45 minutes, resuspended in fresh buffer to final concentration ∼ 700 cells μL^−1^, and incubated for another 90 minutes. The assay began with the addition of a 2 nM radio-labeled tracer, followed by 10 µM NPA, with samples taken every 90 seconds. Cells were then treated with 500 µL of 96% (v/v) ethanol for 30 minutes and mixed with scintillation cocktail (EcoLite(+)™, MP Biomedicals, CA, USA). The radioactivity was measured using Tri-Carb 2900TR scintillation counter (PerkinElmer, CT, USA).

### Gravitropic bending assay

Seedlings were grown on vertical plates under cycles of 16h light/8h dark for 4 days. The plates were then turned 90° and incubated vertically for one day. Images were taken after 24h of gravistimulation and directional angle of root tips were measured in Image J software.

### Confocal and brightfield microscopy

*Chara braunii* brightfield images of thallus and antheridia were acquired on Olympus Provis AX-70 (Olympus, Japan) with a color digital camera (Olympus DP28, Japan) using DP2-AOU imaging software. Confocal laser scanning microscopy was performed using Leica SP8 with the HC PL APO CS2 63×/1.2 W objective (GFP, Alexa Fluor 488: λex=488 nm, λem=490-550, Alexa Fluor 546: λex=546, λem=570-630, FM 4-64: λex=488 nm, λem= >590 nm, DAPI, calcoflur and Hoechst 33342 λex= 405, λem=390-490 nm), the pinhole was set to 1 AU at 580 nm. The equatorial sections of BY-2 cells estradiol induced 5-d-old BY-2 cells expressing XVE:CbPINa:GFP or XVE:CbPINc:GFP were acquired by Leica SP8 confocal microscope (HC PL APO CS2 63×/1.2 W objective, image resolution ∼5 px µm^−1^) with the same detection range as for *Chara braunii*. The confocal images of *Arabidopsis thaliana* were detected using confocal microscope (Zeiss, LSM 800, inverted). For imaging GFP and FM4-64, the 488 nm laser (GFP:λem=500-550, FM 4-64: λem= 670-740 nm) was used. The confocal images of *Marchantia polymorpha* were taken with a Leica SP5 upright confocal microscope (GFP: λex=488 nm, λem=505-515). The objective used was a 10x NA 0.30 dipping lens (HCX APO L U-V-I 10x/0.30); the agar plate growing 1-day-old gemmalings was filled with water immediately before imaging. Gametangiophore stalks were cut down the center of the stalk with a razor blade before mounting to a plate lid with Blu-Tack which was filled with water for imaging. For white light images of Marchantia gametangiophore stalks, a SteREOLumar v.12 dissecting microscope, 90.63 FWD 81 mm PlanApo S objective with camera AxioCamHRc (Zeiss), was used. All Images were processed using Fiji (ImageJ) by creating z-stacks, selecting channels colors under “Lookup Tables”, and adjusting “Brightness/Contrast” settings.

### *In silico* molecular docking

Protein structures were predicted using ColabFold (Mirdita et al. 2022), which integrates multi-sequence alignment with AlphaFold (Jumper et al. 2021). The structures were relaxed in water using the CHARMM36 force field (Best et al. 2012) and GROMACS (Abraham et al. 2015) and visualized with ChimeraX (Goddard et al. 2018). Ligands were docked using AutoDockFR (Zhao et al. 2006; Ravindranath et al. 2015), initially placed based on IAA or NPA poses in *AtPIN8* (Ung et al. 2022). Affinity grids were automatically set with 4.0 Å padding and 0.375 Å spacing. Each docking included 50 runs with 25 million evaluations, and results were visualized using LigPlot+ (Laskowski and Swindells 2011).

### Multiple sequence alignment

The Arabidopsis PIN1, Marchantia PIN1 and *Chara braunii* PINa and c were aligned in Geneious Prime 2021.2.2 using MUSCLE 3.8.425 (Edgar 2004) algorithm.

### Quantification and statistical analysis

#### *Chara braunii* regeneration experiments

The thallus length and side branch numbers were measured after 15 days and analyzed using GLMM in the lme4 package (Bates et al. 2015). Gamma and Poisson distributions were used for shoot lengths and numbers, respectively. Three experiment replications were treated as a random effect. Treatment significance was assessed by likelihood-ratio test, and differences between treatments were analyzed using 95% confidence intervals with the emmeans package (Searle et al. 1980).

### Analysis of cytoplasmic streaming

From full-length movies, 2-s long samples were excised each 2-30 s (random intervals) and converted to TIFFs in FFmpeg. Kymograms in manually defined outlines of the longest cell were acquired by KymographClear (Mangeol et al. 2016) macro in Fiji. Individual trajectories were identified by AI-based KymoButler tool (Jakobs et al. 2019), fitted by 2-segments line. The effect of treatment was tested by nonparametric bootstrapping in R.

### Quantification of plasma membrane fluorescent signal

Plasma membrane affinity of BY-2 cells expressing XVE::CbPINa:GFP and XVE::CbPINc:GFP was quantified by a single-dimensional deconvolution-based method (Vosolsobě et al. 2017).

### Quantification of gravitropic bending assay

For each variant, length and direction of the resultant vector were calculated and tested by non-parametric bootstrapping in comparison to a random set of directions.

### Statistical analysis of auxin transport assays

The whole experiment with orthogonal design of induction and NPA treatment was done in triplicates and repeated two times. Asymptotic growth curves (SSasymp) were fitted by nls to individual replications and dependency of predicted asymptotic parameters to estradiol and NPA treatments were fitted by lme4::lmer and tested by χ2-test. Individual differences were tested by emmeans::pairs function.

### RNA-seq quantification

Publicly available transcriptomic libraries (Source data for Fig. S4b) were quantified by two independent approaches, firstly, Kallisto 0.48.0 with *C. braunii* CDS from genome GCA_003427395 as a reference was used, and secondly, reads were aligned to the genomic reference GCA_003427395 by 2-pass mapping in STAR 2.7.10 and next quantified in RSEM 1.3.3 with option for strand-specific protocol, if the ratio of reads mapper to sense and antisense strain was greater than 10 in STAR quantification.

## Acknowledgements

The authors would like to thank Ilse Foissner and Margit Höftberger for discussing details of immunostaining protocol, Katarzyna Retzer and Jan Martinek for help with Western blots, Anna Kampová for help with phosphoproteome sampling, Anja Holzhausen and MadLAnd for providing *Chara braunii* strain S276, and Roman Skokan for valuable discussion.

## Author contributions

K.K., S.V., and J.P. conceptualized the study and provided funding. K.K. performed *Chara braunii* experiments that included cultivation, harvesting, immunolocalization, and growth responses, cloned *Chara* PINs, transformed BY2, and performed Western blot. P.P. performed immunolocalization of tubulin in *Chara*. S.V. computed RNA-seq results, performed statistical analysis of growth experiments, cytoplasmic streaming and STRING analysis. K.M isolated RNA from *Chara* and performed RT-PCR. D.N. performed auxin accumulation assays and *in silico* PIN docking. P.D. measured IAA metabolism. V.S. analyzed metabolism data. A.K. performed isolation of proteins for phosphoproteomics, phosphoenrichment, and phosphoproteomic analysis. A.S. performed *Arabidopsis thaliana* experiments. T.F. performed *Marchatia polymorpha* experiments. D.W., J.F., J.B., and J.P. revised the manuscript.

## Data and code availability

Complete source codes are deposited at https://github.com/vosolsob/chara.

## References

Abraham MJ, Murtola T, Schulz R, Páll S, Smith JC, Hess B, and Lindahl E. GROMACS: High performance molecular simulations through multi-level parallelism from laptops to supercomputers. SoftwareX. 2015:1–2:19–25. 10.1016/j.softx.2015.06.001

Adamowski M and Friml J. PIN-Dependent Auxin Transport: Action, Regulation, and Evolution. The Plant Cell. 2015:27(1):20–32. 10.1105/tpc.114.134874

An G. High Efficiency Transformation of Cultured Tobacco Cells 1. Plant Physiol. 1985:79(2):568–570.

Bates D, Mächler M, Bolker B, and Walker S. Fitting Linear Mixed-Effects Models Using lme4. Journal of Statistical Software. 2015:67:1–48. 10.18637/jss.v067.i01

Becker B and Marin B. Streptophyte algae and the origin of embryophytes. Ann Bot. 2009:103(7):999–1004. 10.1093/aob/mcp044

Beilby MJ, Turi CE, Baker TC, Tymm FJ, and Murch SJ. Circadian changes in endogenous concentrations of indole-3-acetic acid, melatonin, serotonin, abscisic acid and jasmonic acid in Characeae (*Chara australis* Brown). Plant Signaling & Behavior. 2015:10(11):e1082697. 10.1080/15592324.2015.1082697

Bennett T, Brockington SF, Rothfels C, Graham SW, Stevenson D, Kutchan T, Rolf M, Thomas P, Wong GK-S, Leyser O, et al. Paralogous Radiations of PIN Proteins with Multiple Origins of Noncanonical PIN Structure. Molecular Biology and Evolution. 2014:31(8):2042–2060. 10.1093/molbev/msu147

Best RB, Zhu X, Shim J, Lopes PEM, Mittal J, Feig M, and MacKerell AD Jr. Optimization of the Additive CHARMM All-Atom Protein Force Field Targeting Improved Sampling of the Backbone ϕ, ψ and Side-Chain χ1 and χ2 Dihedral Angles. J Chem Theory Comput. 2012:8(9):3257–3273. 10.1021/ct300400x

Bierenbroodspot MJ, Pröschold T, Fürst-Jansen JMR, de Vries S, Irisarri I, Darienko T, and de Vries J. Phylogeny and evolution of streptophyte algae. Annals of Botany. 2024:134(3):385–400. 10.1093/aob/mcae091

Blake TJ, Reid DM, and Rood SB. Ethylene, indoleacetic acid and apical dominance in peas: A reappraisal. Physiologia Plantarum. 1983:59(3):481–487. 10.1111/j.1399-3054.1983.tb04234.x

Boot KJM, Libbenga KR, Hille SC, Offringa R, and van Duijn B. Polar auxin transport: an early invention. Journal of Experimental Botany. 2012:63(11):4213–4218. 10.1093/jxb/ers106

Bowman JL. The origin of a land flora. Nat Plants. 2022:8(12):1352–1369. 10.1038/s41477-022-01283-y

Bowman JL, Flores Sandoval E, and Kato H. On the Evolutionary Origins of Land Plant Auxin Biology. Cold Spring Harb Perspect Biol. 2021:13(6):a040048. 10.1101/cshperspect.a040048

Bowman JL, Kohchi T, Yamato KT, Jenkins J, Shu S, Ishizaki K, Yamaoka S, Nishihama R, Nakamura Y, Berger F, et al. Insights into Land Plant Evolution Garnered from the Marchantia polymorpha Genome. Cell. 2017:171(2):287–304.e15. 10.1016/j.cell.2017.09.030

Buschmann H. Into another dimension: how streptophyte algae gained morphological complexity. Journal of Experimental Botany. 2020:71(11):3279–3286. 10.1093/jxb/eraa181

Carrillo-Carrasco VP, Hernandez-Garcia J, Mutte SK, and Weijers D. The birth of a giant: evolutionary insights into the origin of auxin responses in plants. The EMBO Journal. 2023:42(6):e113018. 10.15252/embj.2022113018

Causse HB. 1993. Marchantia L: the European and African Taxa (J Cramer: Berlin).

Chau R, Bisson MA, Siegel A, Elkin G, Klim P, and Straubinger RM. Distribution of Charasomes in Chara: Re-establishment and Loss in Darkness and Correlation with Banding and Inorganic Carbon Uptake. Functional Plant Biol. 1994:21(1):113–123. 10.1071/pp9940113

Cheng S, Xian W, Fu Y, Marin B, Keller J, Wu T, Sun W, Li X, Xu Y, Zhang Y, et al. Genomes of Subaerial Zygnematophyceae Provide Insights into Land Plant Evolution. Cell. 2019:179(5):1057–1067.e14. 10.1016/j.cell.2019.10.019

Clabeaux BL. Potential use of Charophytes in the Phytoremediation of Cadmium Contaminated Soils. 2011. Doctoral thesis. University at Buffalo.

Delbarre A, Muller P, Imhoff V, and Guern J. Comparison of mechanisms controlling uptake and accumulation of 2,4-dichlorophenoxy acetic acid, naphthalene-1-acetic acid, and indole-3-acetic acid in suspension-cultured tobacco cells. Planta. 1996:198(4):532–541. 10.1007/BF00262639

Dibb-Fuller JE and Morris DA. Studies on the evolution of auxin carriers and phytotropin receptors: Transmembrane auxin transport in unicellular and multicellular Chlorophyta. Planta. 1992:186(2):219–226. 10.1007/BF00196251

Donoghue PCJ, Harrison CJ, Paps J, and Schneider H. The evolutionary emergence of land plants. Current Biology. 2021:31(19):R1281–R1298. 10.1016/j.cub.2021.07.038

Feng X, Zheng J, Irisarri I, Yu H, Zheng B, Ali Z, de Vries S, Keller J, Fürst-Jansen JMR, Dadras A, et al. Genomes of multicellular algal sisters to land plants illuminate signaling network evolution. Nat Genet. 2024:56(5):1018–1031. 10.1038/s41588-024-01737-3

Fisher TJ, Flores-Sandoval E, Alvarez JP, and Bowman JL. PIN-FORMED is required for shoot phototropism/gravitropism and facilitates meristem formation in Marchantia polymorpha. New Phytologist. 2023:238(4):1498–1515. 10.1111/nph.18854

Flores-Sandoval E, Eklund DM, and Bowman JL. A Simple Auxin Transcriptional Response System Regulates Multiple Morphogenetic Processes in the Liverwort Marchantia polymorpha. PLOS Genetics. 2015:11(5):e1005207. 10.1371/journal.pgen.1005207

Foissner I, Sommer A, and Hoeftberger M. Photosynthesis-dependent formation of convoluted plasma membrane domains in Chara internodal cells is independent of chloroplast position. Protoplasma. 2015:252(4):1085–1096. 10.1007/s00709-014-0742-9

Friml J, Gallei M, Gelová Z, Johnson A, Mazur E, Monzer A, Rodriguez L, Roosjen M, Verstraeten I, Živanović BD, et al. ABP1-TMK auxin perception for global phosphorylation and auxin canalization. Nature. 2022:609(7927):575–581. 10.1038/s41586-022-05187-x

Gamborg OL, Miller RA, and Ojima K. Nutrient requirements of suspension cultures of soybean root cells. Experimental Cell Research. 1968:50(1):151–158. 10.1016/0014-4827(68)90403-5

Gao X, Nagawa S, Wang G, and Yang Z. Cell Polarity Signaling: Focus on Polar Auxin Transport. Molecular Plant. 2008:1(6):899–909. 10.1093/mp/ssn069

Goddard TD, Huang CC, Meng EC, Pettersen EF, Couch GS, Morris JH, and Ferrin TE. UCSF ChimeraX: Meeting modern challenges in visualization and analysis. Protein Science. 2018:27(1):14–25. 10.1002/pro.3235

Godlewski M. Auxin as a factor regulating initiation, of mitosis in antheridial filaments of *Chara* vulgaris L. Biochemie und Physiologie der Pflanzen. 1980:175(4):314–321. 10.1016/S0015-3796(80)80072-2

Hackenberg D and Pandey S. Heterotrimeric G-proteins in green algae: an early innovation in the evolution of the plant lineage. Plant Signal Behav. 2014:9(4):e28457. 10.4161/psb.28457

Hallgren J, Tsirigos KD, Pedersen MD, Armenteros JJA, Marcatili P, Nielsen H, Krogh A, and Winther O. DeepTMHMM predicts alpha and beta transmembrane proteins using deep neural networks. 2022:2022.04.08.487609. 10.1101/2022.04.08.487609

Harrison J. Development and genetics in the evolution of land plant body plans. Philosophical Transactions of the Royal Society B: Biological Sciences. 2017:372(1713):20150490. 10.1098/rstb.2015.0490

Hernández-García J, Carrillo-Carrasco VP, Rienstra J, Tanaka K, Roij M de, Dipp-Álvarez M, Freire-Ríos A, Crespo I, Boer DR, Berg WAM van den, et al. Evolutionary Origins and Functional Diversification of Auxin Response Factors. 2024:2024.08.14.607941. 10.1101/2024.08.14.607941

Heß D, Holzhausen A, and Hess WR. Insight into the nodal cells transcriptome of the streptophyte green alga *Chara* braunii S276. Physiologia Plantarum. 2023:175(5):e14025. 10.1111/ppl.14025

Holzhausen A, Stingl N, Rieth S, Kühn C, Schubert H, and Rensing SA. Establishment and optimization of a new model organism to study early land plant evolution: germination, cultivation and oospore variation of *Chara braunii* Gmelin, 1826. Front Plant Sci. 2022:13(987741):987741. 10.3389/fpls.2022.987741

Hori K, Maruyama F, Fujisawa T, Togashi T, Yamamoto N, Seo M, Sato S, Yamada T, Mori H, Tajima N, et al. Klebsormidium flaccidum genome reveals primary factors for plant terrestrial adaptation. Nat Commun. 2014:5(1):3978. 10.1038/ncomms4978

Ishizaki K, Chiyoda S, Yamato KT, and Kohchi T. Agrobacterium-Mediated Transformation of the Haploid Liverwort Marchantia polymorpha L., an Emerging Model for Plant Biology. Plant and Cell Physiology. 2008:49(7):1084–1091. 10.1093/pcp/pcn085

Ishizaki K, Nishihama R, Ueda M, Inoue K, Ishida S, Nishimura Y, Shikanai T, and Kohchi T. Development of Gateway Binary Vector Series with Four Different Selection Markers for the Liverwort Marchantia polymorpha. PLOS ONE. 2015:10(9):e0138876. 10.1371/journal.pone.0138876

Jahnke E and Libbert E. Indol-3-essigsäure als einziges natives Auxin bei *Chara*-Arten. Z Bot. 1964:52:283–290.

Jakobs MA, Dimitracopoulos A, and Franze K. KymoButler, a deep learning software for automated kymograph analysis. eLife. 2019:8:e42288. 10.7554/eLife.42288

Jin Q, Scherp P, Heimann K, and Hasenstein KH. Auxin and cytoskeletal organization in algae. Cell Biol Int. 2008:32(5):542–545. 10.1016/j.cellbi.2007.11.005

Jumper J, Evans R, Pritzel A, Green T, Figurnov M, Ronneberger O, Tunyasuvunakool K, Bates R, Žídek A, Potapenko A, et al. Highly accurate protein structure prediction with AlphaFold. Nature. 2021:596(7873):583–589. 10.1038/s41586-021-03819-2

Klämbt D, Knauth B, and Dittmann I. Auxin dependent growth of rhizoids of *Chara* globularis. Physiologia Plantarum. 1992:85(3):537–540. 10.1111/j.1399-3054.1992.tb05823.x

Kuhn A, Roosjen M, Mutte S, Dubey SM, Carrillo Carrasco VP, Boeren S, Monzer A, Koehorst J, Kohchi T, Nishihama R, et al. RAF-like protein kinases mediate a deeply conserved, rapid auxin response. Cell. 2024:187(1):130–148.e17. 10.1016/j.cell.2023.11.021

Kurtović K, Schmidt V, Nehasilová M, Vosolsobě S, and Petrášek J. Rediscovering *Chara* as a model organism for molecular and evo-devo studies. Protoplasma. 2024:261(2):183–196. 10.1007/s00709-023-01900-3

Laskowski RA and Swindells MB. LigPlot+: Multiple Ligand–Protein Interaction Diagrams for Drug Discovery. J Chem Inf Model. 2011:51(10):2778–2786. 10.1021/ci200227u

Liang Z, Geng Y, Ji C, Du H, Wong CE, Zhang Q, Zhang Y, Zhang P, Riaz A, Chachar S, et al. Mesostigma viride Genome and Transcriptome Provide Insights into the Origin and Evolution of Streptophyta. Advanced Science. 2020:7(1):1901850. 10.1002/advs.201901850

Lv S, Wang L, Zhang X, Li X, Fan L, Xu Y, Zhao Y, Xie H, Sawchuk M, Scarpella E, et al. Arabidopsis NHX5 and NHX6 regulate PIN6-mediated auxin homeostasis and growth. Journal of Plant Physiology. 2020:255:153305. 10.1016/j.jplph.2020.153305

Mangeol P, Prevo B, and Peterman EJG. KymographClear and KymographDirect: two tools for the automated quantitative analysis of molecular and cellular dynamics using kymographs. MBoC. 2016:27(12):1948–1957. 10.1091/mbc.e15-06-0404

Mirdita M, Schütze K, Moriwaki Y, Heo L, Ovchinnikov S, and Steinegger M. ColabFold: making protein folding accessible to all. Nat Methods. 2022:19(6):679–682. 10.1038/s41592-022-01488-1

Müller K, Hošek P, Laňková M, Vosolsobě S, Malínská K, Čarná M, Fílová M, Dobrev PI, Helusová M, Hoyerová K, et al. Transcription of specific auxin efflux and influx carriers drives auxin homeostasis in tobacco cells. The Plant Journal. 2019:100(3):627–640. 10.1111/tpj.14474

Murashige T and Skoog F. A Revised Medium for Rapid Growth and Bio Assays with Tobacco Tissue Cultures. Physiologia Plantarum. 1962:15(3):473–497. 10.1111/j.1399-3054.1962.tb08052.x

Nishiyama T, Sakayama H, de Vries J, Buschmann H, Saint-Marcoux D, Ullrich KK, Haas FB, Vanderstraeten L, Becker D, Lang D, et al. The *Chara* genome: secondary complexity and implications for plant terrestrialization. Cell. 2018:174(2):448–464. 10.1016/j.cell.2018.06.033

Mutte SK, Kato H, Rothfels C, Melkonian M, Wong GK-S, and Weijers D. Origin and evolution of the nuclear auxin response system. eLife. 2018:7:e33399. 10.7554/eLife.33399

Naramoto S. Polar transport in plants mediated by membrane transporters: focus on mechanisms of polar auxin transport. Current Opinion in Plant Biology. 2017:40:8–14. 10.1016/j.pbi.2017.06.012

Petrášek J, Černá A, Schwarzerová K, Elčkner M, Morris DA, and Zažímalová E. Do Phytotropins Inhibit Auxin Efflux by Impairing Vesicle Traffic? Plant Physiol. 2003:131(1):254–263. 10.1104/pp.012740

Pan L, Fonseca De Lima CF, Vu LD, and De Smet I. A Comprehensive Phylogenetic Analysis of the MAP4K Family in the Green Lineage. Front Plant Sci. 2021:12. 10.3389/fpls.2021.650171

Petrášek J, Mravec J, Bouchard R, Blakeslee JJ, Abas M, Seifertová D, Wiśniewska J, Tadele Z, Kubeš M, Čovanová M, et al. PIN Proteins Perform a Rate-Limiting Function in Cellular Auxin Efflux. Science. 2006:312(5775):914–918. 10.1126/science.1123542

Pollak B, Cerda A, Delmans M, Álamos S, Moyano T, West A, Gutiérrez RA, Patron NJ, Federici F and Haseloff J. Loop assembly: a simple and open system for recursive fabrication of DNA circuits. New Phytol. 2019:222: 628–640. 10.1111/nph.15625

Ravindranath PA, Forli S, Goodsell DS, Olson AJ, and Sanner MF. AutoDockFR: Advances in Protein-Ligand Docking with Explicitly Specified Binding Site Flexibility. PLOS Computational Biology. 2015:11(12):e1004586. 10.1371/journal.pcbi.1004586

Sauer M and Kleine-Vehn J. PIN-FORMED and PIN-LIKES auxin transport facilitators. Development. 2019:146(15):dev168088. 10.1242/dev.168088

Schmidt V, Skokan R, Depaepe T, Kurtović K, Haluška S, Vosolsobě S, Vaculíková R, Pil A, Dobrev PI, Motyka V, et al. Phytohormone profiling in an evolutionary framework. Nat Commun. 2024:15(1):3875. 10.1038/s41467-024-47753-z

Schmölzer PM, Höftberger M, and Foissner I. Plasma membrane domains participate in pH banding of Chara internodal cells. Plant Cell Physiol. 2011:52(8):1274–1288. 10.1093/pcp/pcr074

Searle SR, Speed FM, and Milliken GA. Population Marginal Means in the Linear Model: An Alternative to Least Squares Means. The American Statistician. 1980:34(4):216–221. 10.1080/00031305.1980.10483031

Skokan R, Medvecká E, Viaene T, Vosolsobě S, Zwiewka M, Müller K, Skůpa P, Karady M, Zhang Y, Janacek DP, et al. PIN-driven auxin transport emerged early in streptophyte evolution. Nat Plants. 2019:5(11):1114–1119. 10.1038/s41477-019-0542-5

Su L, Zhang T, Yang B, Dong T, Liu X, Bai Y, Liu H, Xiong J, Zhong Y, and Cheng Z-M (Max). Different evolutionary patterns of TIR1/AFBs and AUX/IAAs and their implications for the morphogenesis of land plants. BMC Plant Biol. 2023:23:265. 10.1186/s12870-023-04253-4

Su N, Zhu A, Tao X, Ding ZJ, Chang S, Ye F, Zhang Y, Zhao C, Chen Q, Wang J, et al. Structures and mechanisms of the Arabidopsis auxin transporter PIN3. Nature. 2022:609(7927):616–621. 10.1038/s41586-022-05142-w

Sztein AE, Cohen JD, and Cooke TJ. Evolutionary Patterns in the Auxin Metabolism of Green Plants. International Journal of Plant Sciences. 2000:161(6):849–859. 10.1086/317566

Ung KL, Schulz L, Stokes DL, Hammes UZ, and Pedersen BP. Substrate recognition and transport mechanism of the PIN-FORMED auxin exporters. Trends in Biochemical Sciences. 2023:48(11):937–948. 10.1016/j.tibs.2023.07.006

Ung KL, Winkler M, Schulz L, Kolb M, Janacek DP, Dedic E, Stokes DL, Hammes UZ, and Pedersen BP. Structures and mechanism of the plant PIN-FORMED auxin transporter. Nature. 2022:609(7927):605–610. 10.1038/s41586-022-04883-y

Vieten A, Vanneste S, Wiśniewska J, Benková E, Benjamins R, Beeckman T, Luschnig C, and Friml J. Functional redundancy of PIN proteins is accompanied by auxin-dependent cross-regulation of PIN expression. Development. 2005:132(20):4521–4531. 10.1242/dev.02027

Vosolsobě S, Petrášek J, and Schwarzerová K. Evolutionary plasticity of plasma membrane interaction in DREPP family proteins. Biochimica et Biophysica Acta (BBA) - Biomembranes. 2017:1859(5):686–697. 10.1016/j.bbamem.2017.01.017

Vosolsobě S, Skokan R, and Petrášek J. The evolutionary origins of auxin transport: what we know and what we need to know. J Exp Bot. 2020:71(11):3287–3295. 10.1093/jxb/eraa169

Wasteneys GO, Jablonsky PP, and Williamson RE. Assembly of purified brain tubulin at cortical and endoplasmic sites in perfused internodal cells of the alga Nitella tasmanica. Cell Biology International Reports. 1989:13(6):513–528. 10.1016/0309-1651(89)90098-2

Weller B, Zourelidou M, Frank L, Barbosa ICR, Fastner A, Richter S, Jürgens G, Hammes UZ, and Schwechheimer C. Dynamic PIN-FORMED auxin efflux carrier phosphorylation at the plasma membrane controls auxin efflux-dependent growth. Proceedings of the National Academy of Sciences. 2017:114(5):E887–E896. 10.1073/pnas.1614380114

Yang Z, Xia J, Hong J, Zhang C, Wei H, Ying W, Sun C, Sun L, Mao Y, Gao Y, et al. Structural insights into auxin recognition and efflux by Arabidopsis PIN1. Nature. 2022:609(7927):611–615. 10.1038/s41586-022-05143-9

Zhang S, de Boer AH, and van Duijn B. Auxin effects on ion transport in Chara corallina. Journal of Plant Physiology. 2016:193:37–44. 10.1016/j.jplph.2016.02.009

Zhang Y, Xiao G, Wang X, Zhang X, and Friml J. Evolution of fast root gravitropism in seed plants. Nat Commun. 2019:10(1):3480. 10.1038/s41467-019-11471-8

Zhao Y, Stoffler D, and Sanner M. Hierarchical and multi-resolution representation of protein flexibility. Bioinformatics. 2006:22(22):2768–2774. 10.1093/bioinformatics/btl481

Zuo J, Niu Q-W, and Chua N-H. An estrogen receptor-based transactivator XVE mediates highly inducible gene expression in transgenic plants. The Plant Journal. 2000:24(2):265–273. 10.1046/j.1365-313x.2000.00868.x

Żabka A, Polit JT, Winnicki K, Paciorek P, Juszczak J, Nowak M, and Maszewski J. PIN2-like proteins may contribute to the regulation of morphogenetic processes during spermatogenesis in Chara vulgaris. Plant Cell Rep. 2016:35(8):1655–1669. 10.1007/s00299-016-1979-x

