## Supplementary information for "The role of IAA and its transport in the complex streptophyte algae *Chara braunii*"

Supplementary information for this article contains:

**Fig. S1** Design of cultivation box for *Chara braunii*.
**Fig. S2** Auxin treatment experiment and plasma membrane staining in axillary branches of *Chara braunii*, strain NIES 1604
**Fig. S3** Concentration of IAA and IAA metabolites in *Chara* biomass and its medium determined by LC/MS
**Fig. S4** Melting curve analysis from RT-qPCR of and expression profiles of *Chara* PINs during life cycle.
**Fig. S5** Negative controls for CbPINa and CbPINc immunostainings
**Fig. S6** Positive controls for CbPINa and CbPINc immunostainings
**Fig. S7** Multiple sequence alignment showing conserved sites between *Arabidopsis thaliana*, *Marchantia polymorpha*, and *Chara braunii* PINa and PINc
**Fig. S8** Auxin transport assays in tobacco BY-2 cells
**Fig. S9** STRING analysis of significantly phosphorylated candidates upon IAA treatment
**Table S1** PIN-FORMED auxin efflux carriers in *Chara braunii*
**Table S2** Ligand biding affinities of CbPINs with IAA, 1-NAA, and NPA calculated in Autodock
**Table S3** Significantly phosphorylated proteins under IAA treatment compared to DMSO
**Table S4** Significantly dephosphorylated proteins under IAA treatment compared to DMSO
**Methods S1** Construction of cultivation box for *Chara braunii*
**Methods S2** Immunolocalization of internodal cells with CbPINs and $H^+$ATPase
**Methods S3** Immunolocalization of antheridia CbPINs and $H^+$ATPase
**Methods S4** Immunolocalization of tubulin in internodal cells

**Methods S5** Immunolocalization of tubulin in antheridial cells

**Methods S6** Protein extraction and Western blot

**Methods S7** Sample preparation for phosphoproteomic analysis

**Methods S8** Protein extraction for phosphoproteomic analysis

**Methods S9** Phosphopeptide enrichment

**Methods S10** Mass spectrometry

**Methods S11** Phosphoproteomic data analysis

**Video S1-16** Cytoplasmic streaming videos (attached as separate files)

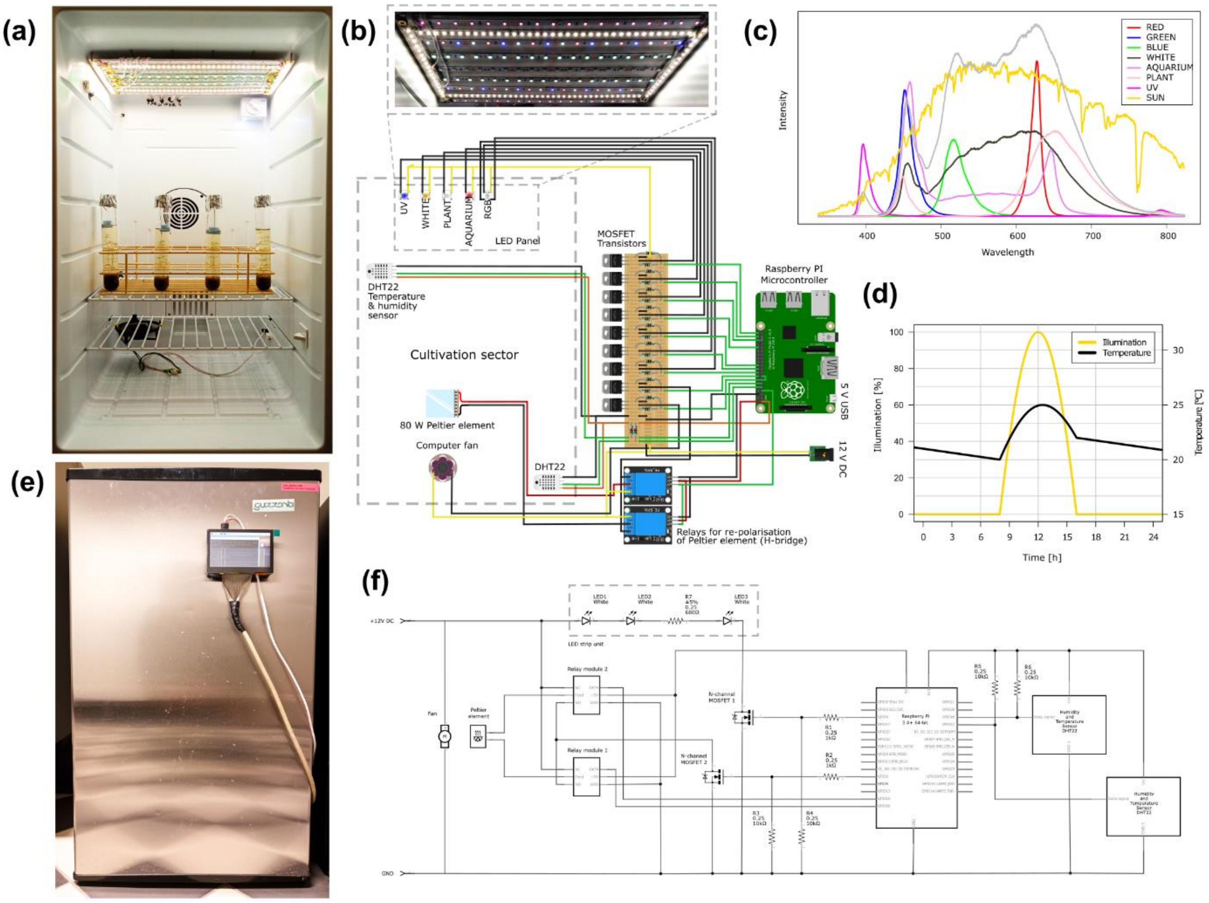


**Fig. S1 Design of cultivation box for *Chara braunii*. (a,e)** Commercial thermoelectric refrigerator is equipped with a custom LED panel **(b)** consisting of RGB, day white, and UV LED strips, as well as special LED strips designed for aquariums and plant growth with extended far-red emission. Below LED strips is shown regulation of one type of LED stripe and regulation of the Peltier element via H-bridge and MOSFET, which is constructed done with respect to the actual temperature measured by a pair of DHT22 sensors. Built Built-in PC fan ensures homogenization of the temperature inside the box. **(c)** The sun spectrum is shown for comparison. The gray line shows an example of the LEDs' intensity setup with maximal similarity to natural sunlight. **(d)** Exemplary day-courses of illumination and temperature showing non-linear simulation of those environmental factors, close to natural situation **(e)** Both illumination and temperature in the box is are regulated by MOSFET transistors and the Raspberry PI microcomputer placed on the fridge's door together with touch display enabling the checking of internal parameters, which are controlled by a python script. **(f)**. Simplified wiring diagram.

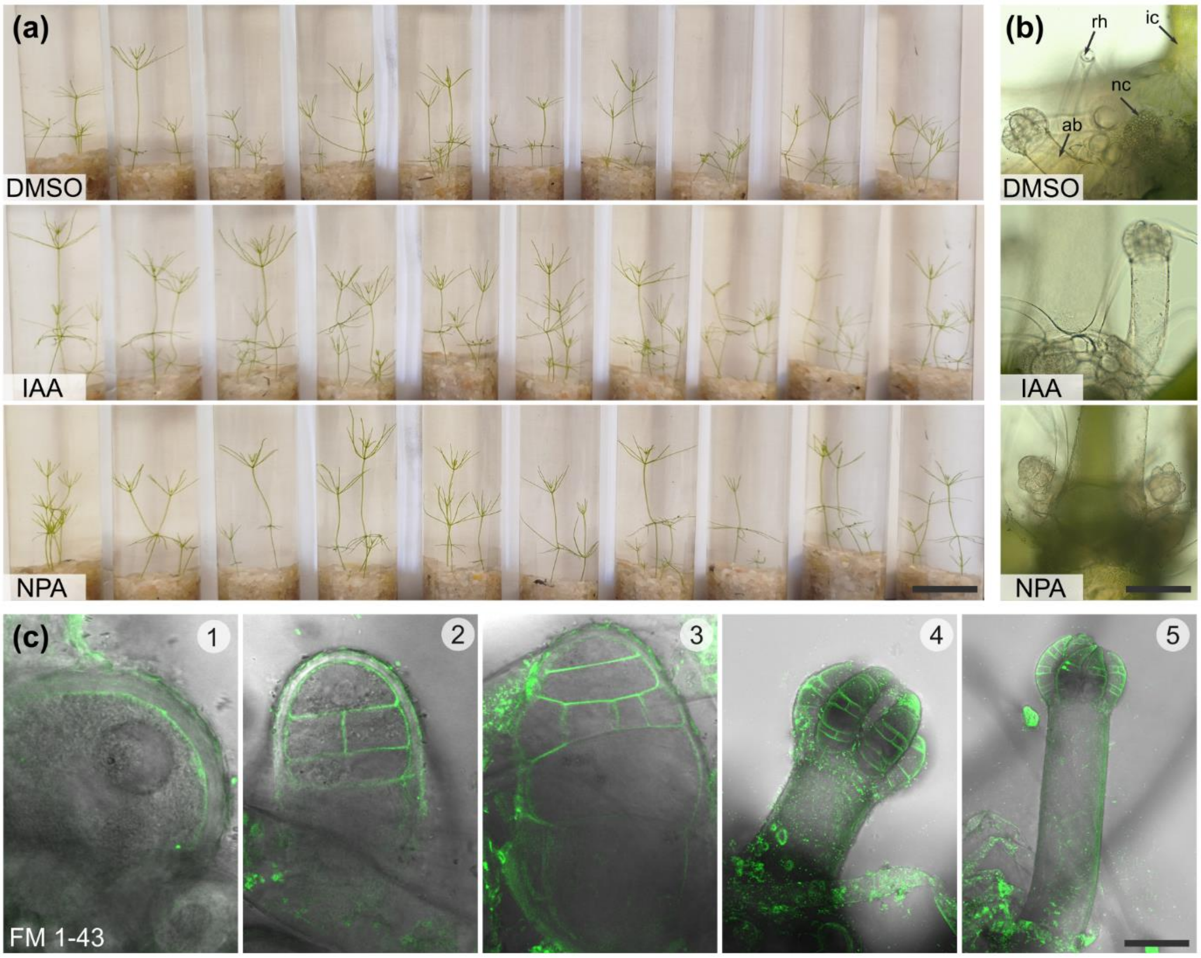


**Fig. S2 Auxin treatment experiment and plasma membrane staining in axillary branches of *Chara braunii*, strain NIES 1604. (a)** Image of a representative thallus regeneration experiment that included treatments with DMSO (control), IAA and NPA The picture was taken at the end of the experiment (day 15). **(b)** Brightfield images of axillary branches at the basal node. The node is underneath the surface of the medium, therefore branches that develop underground are transparent. When they are exposed to light, they start being photosynthetic and develop chloroplasts. Rhizoids (rh), internodal cell (ic), axillary branch (ab), nodal cell (nc). **(c)** FM 1-43 staining of PM in various stages (1-5) of axillary branches that developed underground. Since these organs develop without light, the plasma membrane is smooth compared to the invaginated plasma membrane of internodal cells that form charasomes in response to light.

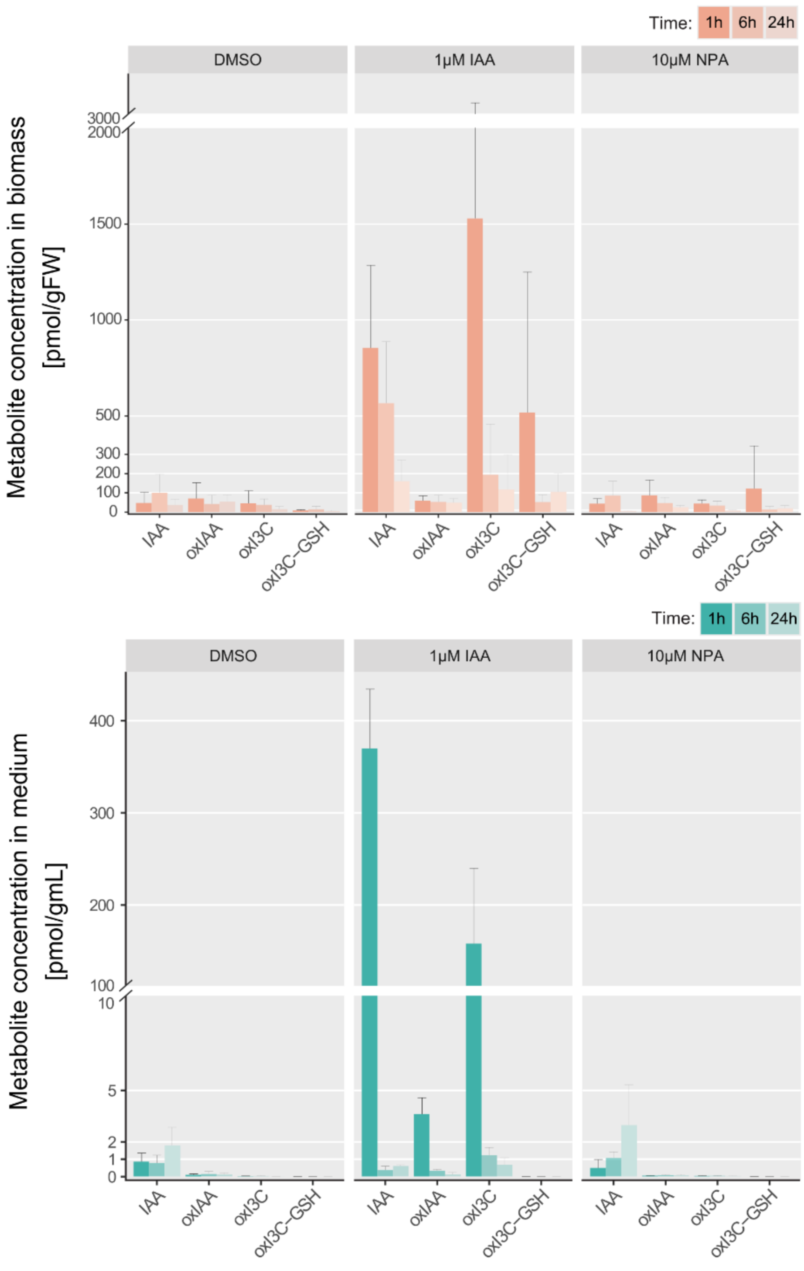


**Fig. S3 Concentration of IAA and IAA metabolites in *Chara* biomass and its medium determined by LC/MS.** The treatments included mock (DMSO), 1 µM IAA and 10 µM NPA. Samples were collected after 1 h, 6 h, and 24 h. 2-oxindole-3-acetic acid (oxIAA), oxindole-3-carbinol (oxI3C), oxindole-3-carbinol-gluthathione (oxI3C-GSH).

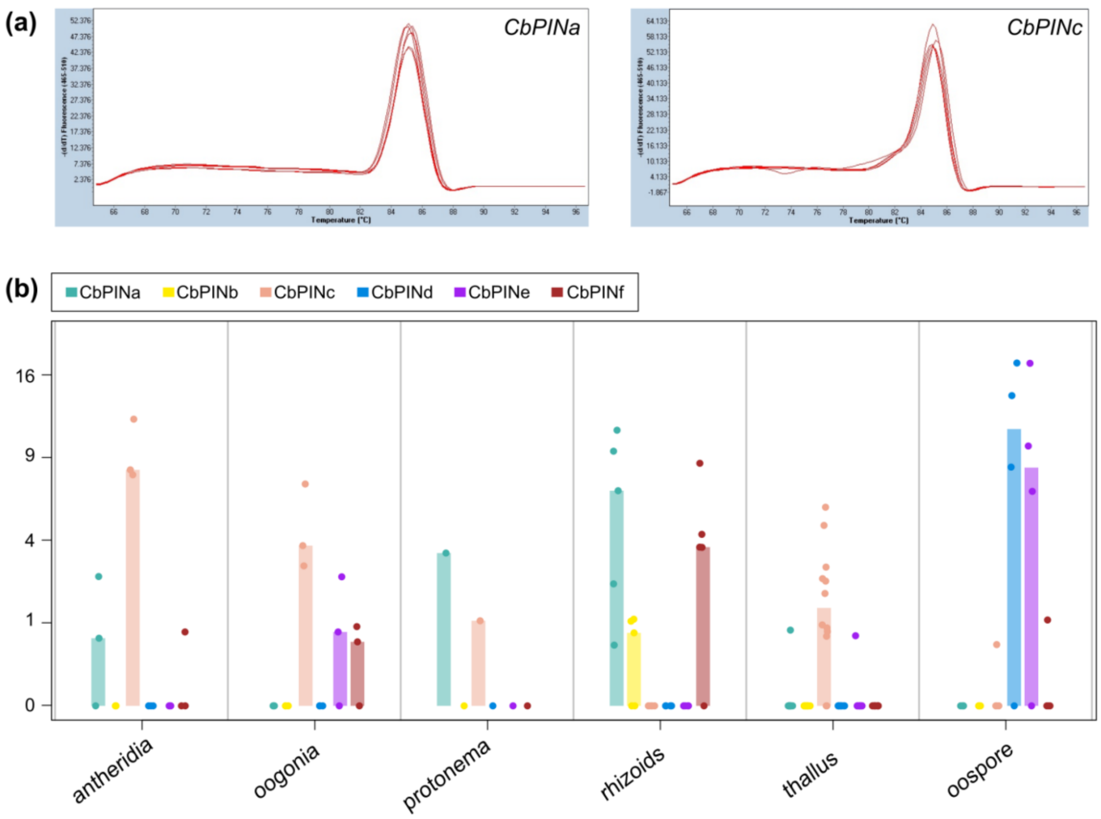


**Fig. S4 Melting curve analysis from RT-qPCR of and expression profiles of *Chara* PINs during the life cycle. (a)** Melt curves from from RT-qPCR of *CbPINa* and *CbPINc*. The dissociation temperatures range from 65°C to 96°C. Amplicons from both *CbPINa* and CbPINc reveal a single peak. **(b)** Data from publicly available RNA-seq libraries (see in Source data) were quantified by Kallisto and STAR. Not**e** higher expressions of *CbPINa* in protonema and rhizoids, while *CbPINc* is expressed in all stages except rhizoids and oospore. The threshold of *TPM* > 0.5 was applied when the results were visualized in R.

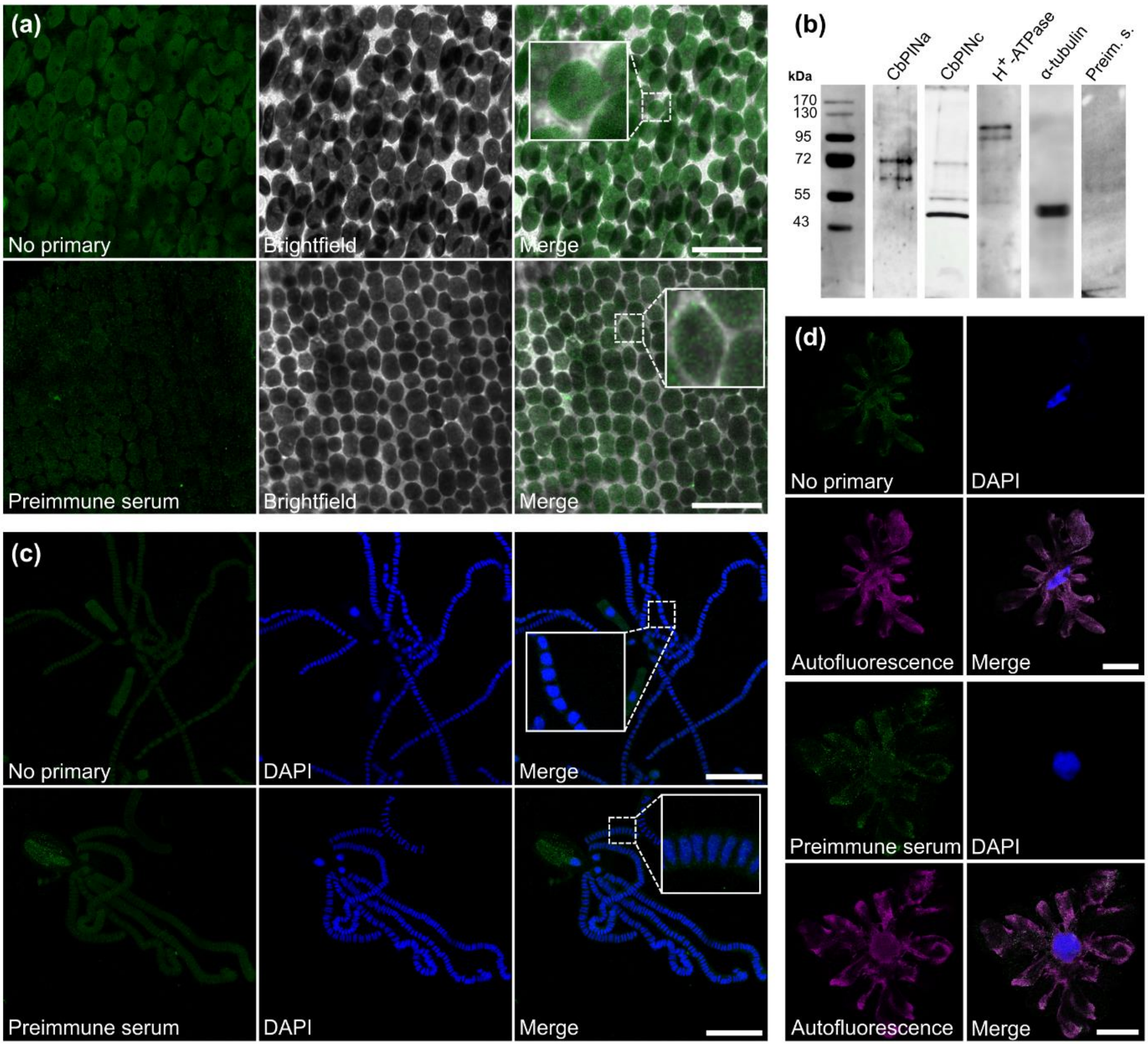


**Fig. S5 Negative controls for CbPINa and CbPINc immunostainings. (a)** Omission of primary antibody and preimmune serum staining in internodal cells. **(b)** Western blots of CbPINa, CbPINc, $H^+$-ATPase, α-tubulin, and preimmune serum. The preimmune serum that served as a negative control shows no visible and on the gel. CbPINa is recognized by a band around 70 kDa which corresponds to the approximate size of the protein, while CbPINc shows 2 bands, one at 70 kDa and the other around 40 kDa which combined corresponds to the CbPINc full length. **(c)** Omission of primary antibody and preimmune serum staining in antheridial filaments. **(d)** Omission of primary antibody, preimmune serum staining, and autofluorescence in antheridia shield cells. Scale bars, 20 µm (a) 50 µm **(b, c, d)**.

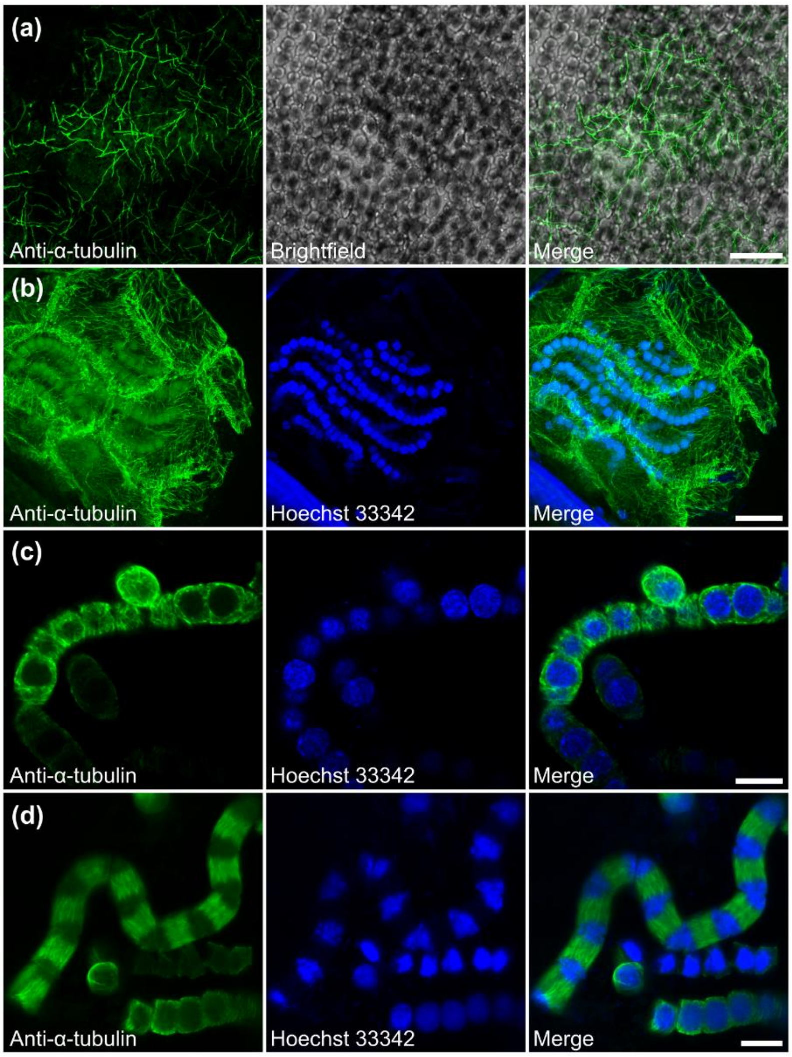


**Fig. S6 Positive controls for CbPINa and CbPINc immunostainings performed by anti-α-tubulin staining. (a)** The internodal cell treated with taxol, confocal, brightfield, and merged image. **(b)** Maximal projection of the whole antheridium. Shield cells are stained with anti-α-tubulin in green, in blue Hoechst stained nuclei of antheridial filaments and merged image. **(c)** Free antheridial filaments showing α-tubulin, Hoechst staining and merged image. **(d)** Antheridial filament during the cell division. Scale bars, 20 µm **(a, b),** 10 µm **(c, d).**

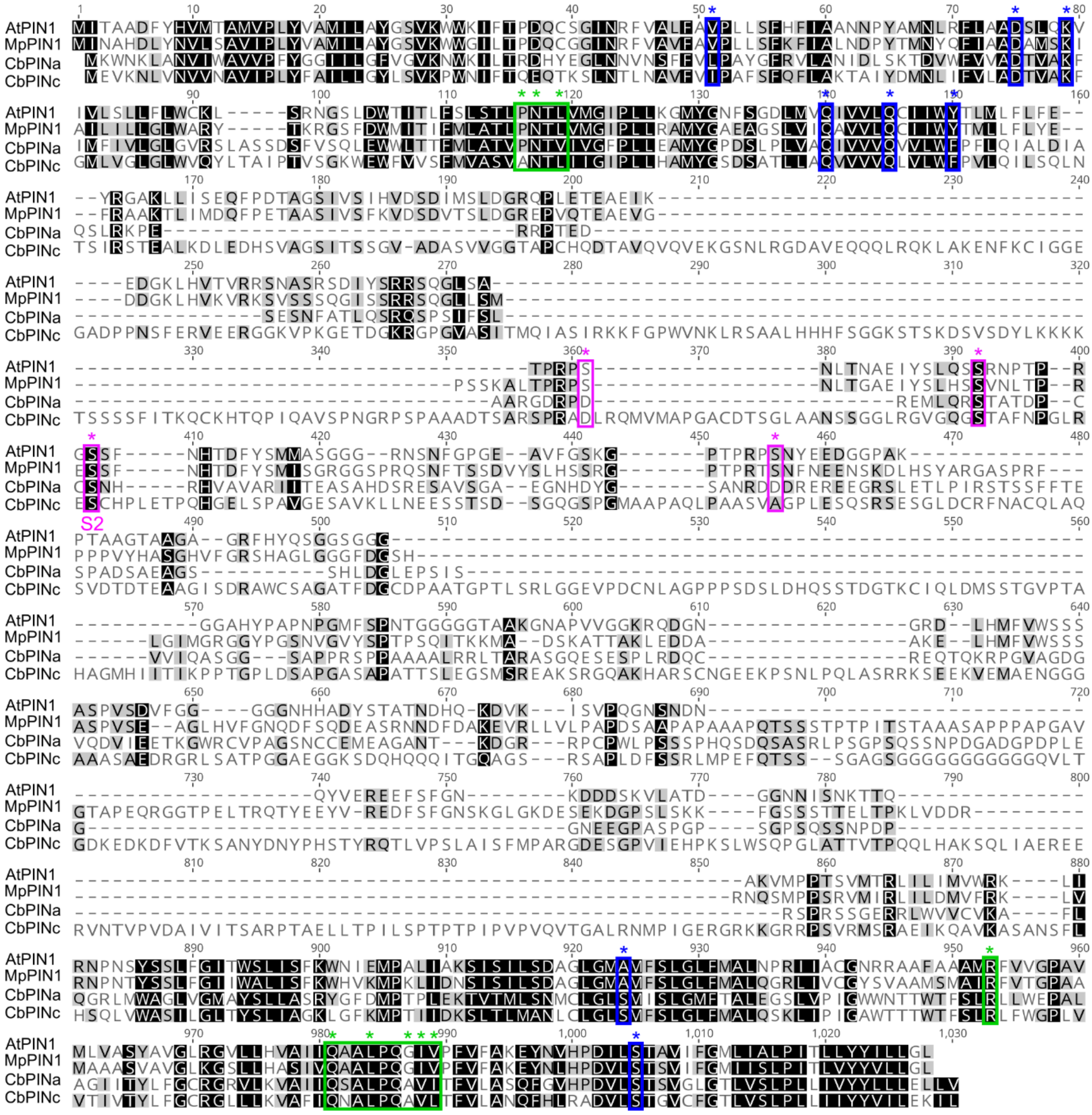

**Fig. S7 Multiple sequence alignment showing conserved sites between *Arabidopsis thaliana*, *Marchantia polymorpha*, and *Chara braunii* PINa and PINc.** The potential phosphorylation sites are framed in magenta. The serine residues that could be potentially phosphorylated are marked with an asterisk, note the S2 phosphorylation site. The green frames represent the sites in TM4 and TM9 that are responsible for the crossover mechanism, and the conserved arginine in TM8 that stabilizes dimerization. Blue frames represent conserved sites that were described to additionally facilitate the auxin transport.

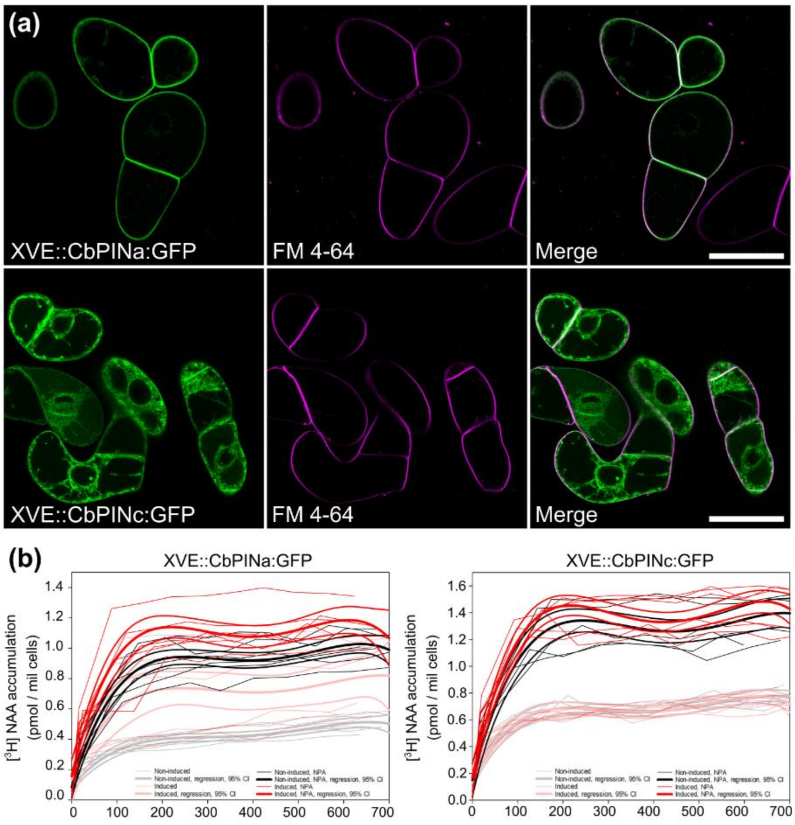


**Fig. S8 Auxin transport assays in tobacco BY-2 cells**. **(a)** Induced tobacco BY-2 cells expressing XVE:CbPINa:GFP and XVE:CbPINc:GFP. The GFP signal at the PM and endoplasmic reticulum, FM 4-64 PM staining and merged images are shown. **(b)** Kinetics of $[^3H]$-NAA accumulation in 5-day-old induced and non-induced XVE:CbPINa:GFP and XVE:CbPINc:GFP BY-2 cells, NPA (10 µM). The thin lines represent independent accumulation runs (*n* = 6, 3 biological repeats, each with two technical repeats), and thick lines are regression lines shown with 95% CI.

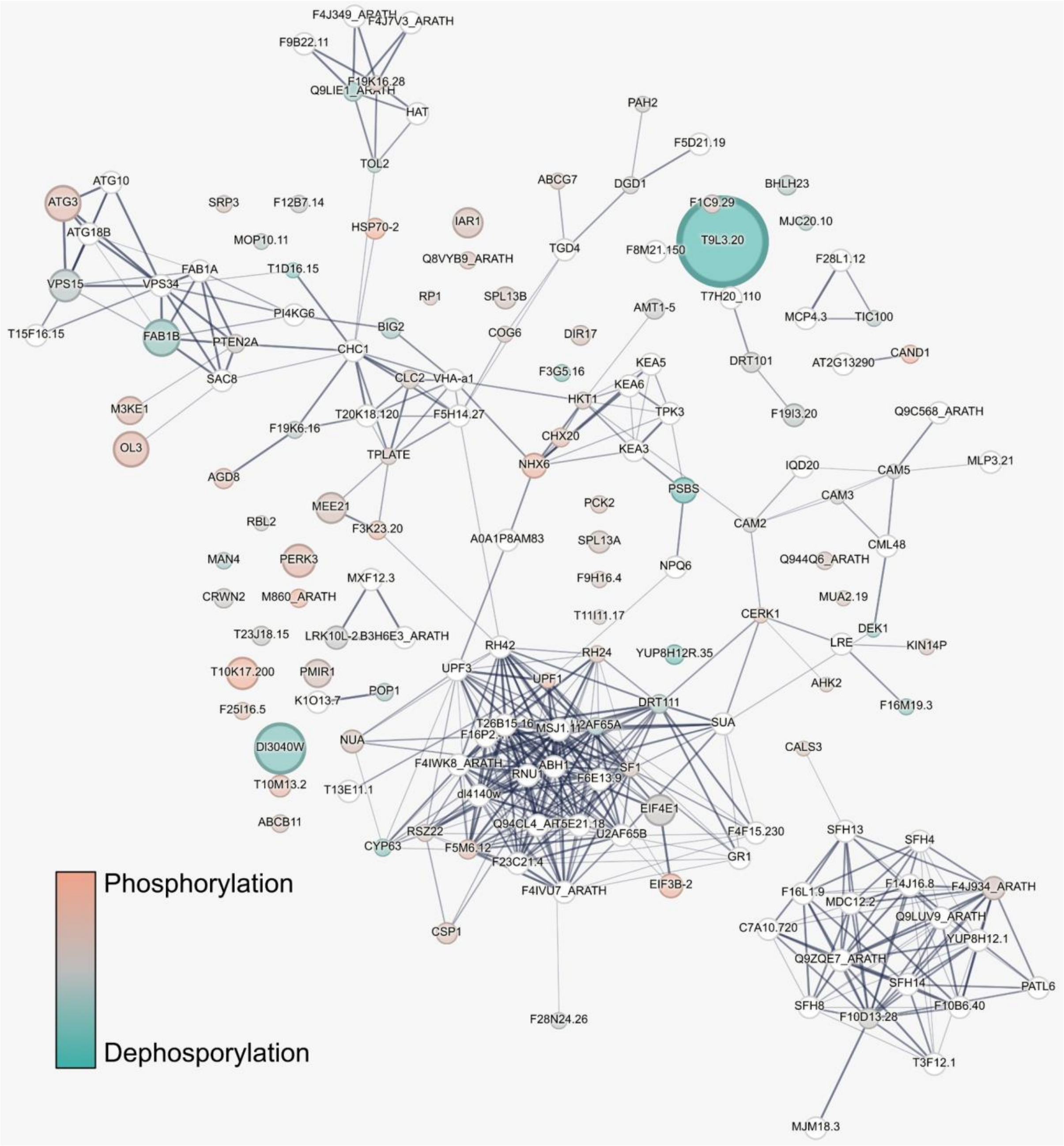


**Fig. S9 STRING analysis of significantly hyper-/dephosphorylated candidates upon 2-min IAA treatment.** Network nodes represent the *A.thaliana* closest homologs to the identified candidates in *Chara braunii*. The color scale is "blue-gray-orange" shows rate of change (phosphorylation/dephosphorylation), the diameter of circles is proportional to significance (-log( *p-value*) of T-test). Second-shell interactors added to the STRING network were shown by empty circles.

**Table S1 PIN-FORMED auxin efflux carriers in *Chara braunii*. Designated PIN protein names and corresponding GenBank and UniProt accessions for protein, gene, and scaffold**

| **Designated name** | **GenBank protein accession** | **UniProt protein accession** | **GenBank gene accession** | **GenBank scaffold** |
|---|---|---|---|---|
| CbPINa* | GBG79698 | A0A388LBL6 | g29962 | BFEA01000325.1 |
| CbPINb | GBG79697 | A0A388LBJ4 | g29961 | BFEA01000325.1 |
| CbPINc | GBG64280 | A0A388K2K4 | g41200 | BFEA01000048.1 |
| CbPINd | GBG90244 | A0A388M6M8 | g50423 | BFEA01000798.1 |
| | GBG90245 | A0A388M6W5 | | |
| CbPINe | GBG90247 | A0A388M6M5 | g50425 | BFEA01000798.1 |
| CbPINf | GBG42734 | A0A388JJV1 | g84230 | BFEA01006036.1 |

*An updated full nucleotide sequence of CbPINa used for this study is deposited in GenBank with accession number PQ421084.

**Table S2 Ligand binding affinities of CbPINs with IAA, 1-NAA and NPA calculated in Autodock**

| **Substrate** | **CbPINa (kcal/mol)** | **CbPINc (kcal/mol)** |
|---|---|---|
| IAA | -7.3 | -6.7 |
| NAA | -7.1 | -5.9 |
| NPA | -7.4 | -6.4 |

**Table S3 Significantly hyperphosphorylated proteins under IAA treatment compared to DMSO.** 30 phosphosites with the highest fold change are shown.

| **UniProt protein accession** | **Protein name** | **Fold Change (Log2)** | **FDR** | **Position** |
|---|---|---|---|---|
| A0A388KUQ6 | Uncharacterized protein | 3,39084 | 1,8647 | 988 |
| A0A388MCN6 | Sodium/hydrogen exchanger | 3,34485 | 3,9272 | 502 |
| A0A388MER3 | DUF4283 domain-containing protein | 3,23836 | 2,8414 | 282 |
| A0A388KL36 | Uncharacterized protein | 3,18321 | 5,1692 | 754 |
| A0A388KWQ6 | Reverse transcriptase domain-containing protein | 3,12867 | 1,6476 | 3374 |
| A0A388JYD9 | Eukaryotic translation initiation factor 3 subunit B | 2,9919 | 2,1406 | 6 |
| A0A388KWF0 | TATA-binding protein interacting (TIP20) domain-containing protein | 2,9870 | 1,6313 | 2 |
| A0A388MCN6 | Sodium/hydrogen exchanger | 2,8504 | 2,2132 | 499 |
| A0A388JT36 | K Homology domain-containing protein | 2,5037 | 1,3836 | 2128 |
| A0A388K1B5 | Helicase ATP-binding domain-containing protein | 2,2454 | 1,3621 | 8 |
| A0A388LES0 | Calponin-homology (CH) domain-containing protein | 2,2139 | 1,6627 | 2288 |
| A0A388KF83 | Nucleotide-diphospho-sugar transferase domain-containing protein | 2,1229 | 1,9603 | 18 |
| A0A388L567 | Oleosin | 1,9048 | 3,1299 | 153 |
| A0A388LJZ4 | C3HC-type domain-containing protein | 1,8777 | 1,4175 | 328 |
| A0A388LJZ4 | C3HC-type domain-containing protein | 1,8699 | 1,4400 | 332 |
| A0A388KCH6 | Protein kinase domain-containing protein | 1,8601 | 2,9262 | 1442 |
| A0A388L7R3 | BUD13 homolog | 1,7689 | 1,7364 | 363 |
| A0A388K496 | Arf-GAP domain-containing protein | 1,7661 | 1,8030 | 374 |
| A0A388LX59 | DCD domain-containing protein | 1,7638 | 1,5402 | 391 |
| A0A388JYD9 | Eukaryotic translation initiation factor 3 subunit B | 1,7616 | 1,4501 | 9 |
| A0A388JYD9 | Eukaryotic translation initiation factor 3 subunit B | 1,7616 | 1,4501 | 13 |
| A0A388M738 | Lysin motif receptor-like kinase (LysM-RLK) | 1,6839 | 1,3852 | 366 |
| A0A388M2N1 | Cation/H+ exchanger domain-containing protein | 1,6712 | 1,7008 | 1443 |
| A0A388L963 | Integrase catalytic domain-containing protein | 1,6210 | 3,1593 | 304 |
| A0A388K0S2 | Protein kinase domain-containing protein | 1,5817 | 2,4890 | 1026 |
| A0A388KWB7 | Glyceraldehyde-3-phosphate dehydrogenase | 1,3011 | 0,4009 | 254 |
| A0A388K9B6 | CCHC-type domain-containing protein | 1,1622 | 1,9016 | 246 |
| A0A388KXT1 | Uncharacterized protein | 1,0705 | 1,5940 | 698 |
| A0A388LPE0 | Uncharacterized protein | 0,9805 | 1,5237 | 138 |
| A0A388LCC7 | Protein kinase domain-containing protein | 0,9516 | 1,3968 | 685 |

**Table S4 Significantly hypophosphorylated proteins under IAA treatment compared to DMSO**. 30 phosphosites with the most negative fold change are shown.

| UniProt protein accession | Protein name | Fold Change (Log2) | FDR | Position |
|---|---|---|---|---|
| A0A388LCC7 | Protein kinase domain-containing protein | -5,2696 | 7,8855 | 793 |
| A0A388LCC7 | Protein kinase domain-containing protein | -4,9838 | 2,1019 | 332 |
| A0A388LT00 | Uncharacterized protein | -4,2944 | 2,2694 | 66 |
| A0A388KW32 | Uncharacterized protein | -4,1707 | 1,7532 | 123 |
| A0A388LLH0 | FYVE-type domain-containing protein | -3,7731 | 1,3609 | 96 |
| A0A388L139 | Methyltransferase small domain-containing protein | -3,7573 | 4,4580 | 39 |
| A0A388KV70 | PPIase cyclophilin-type domain-containing protein | -3,4496 | 1,5472 | 347 |
| A0A388KEQ9 | Uncharacterized protein | -3,4427 | 1,5246 | 1143 |
| A0A388K086 | Calpain catalytic domain-containing protein | -3,2605 | 1,3025 | 1662 |
| A0A388KVW7 | GPI transamidase subunit PIG-U | -3,1982 | 1,4446 | 429 |
| A0A388KH38 | 1-phosphatidylinositol-3-phosphate 5-kinase | -3,004 | 3,2468 | 1919 |
| A0A388L8I5 | Uncharacterized protein | -2,9972 | 1,6645 | 149 |
| A0A388K5K4 | UBX domain-containing protein | -2,6527 | 1,6661 | 269 |
| A0A388KT33 | mRNA decay factor PAT1 domain-containing protein | -2,593 | 2,4643 | 481 |
| A0A388KTB5 | Outer arm dynein light chain 1 | -2,5618 | 3,1413 | 783 |
| A0A388KZ99 | Vacuolar protein sorting-associated protein 11 | -2,5134 | 2,1653 | 420 |
| A0A388KE37 | Uncharacterized protein | -2,4793 | 2,5169 | 2464 |
| A0A388KD78 | Mannan endo-1,4-beta-mannosidase | -2,4221 | 1,4895 | 5 |
| A0A388LPG9 | ORM1-like protein 3 | -2,2788 | 1,3649 | 72 |
| A0A388LVU8 | Uncharacterized protein | -2,2172 | 1,5727 | 328 |
| A0A388JJL4 | GAT domain-containing protein (Fragment) | -2,1995 | 1,3237 | 219 |
| A0A388K510 | DNA-repair protein Xrcc1 N-terminal domain-containing protein | -2,1694 | 1,5528 | 709 |
| A0A388KLS2 | SEC7 domain-containing protein | -2,1459 | 1,8358 | 441 |
| A0A388L3H9 | RRM domain-containing protein | -2,0418 | 1,6903 | 44 |
| A0A388K510 | DNA-repair protein Xrcc1 N-terminal domain-containing protein | -1,894 | 1,4811 | 705 |
| A0A388M724 | Uncharacterized protein (Fragment) | -1,7887 | 1,4207 | 123 |
| A0A388KE37 | Uncharacterized protein | -1,7734 | 1,7555 | 2859 |
| A0A388L2U2 | Integrase catalytic domain-containing protein | -1,7718 | 1,4357 | 1566 |
| A0A388L5N4 | Coatomer subunit beta'-2 | -1,6662 | 1,5293 | 852 |
| A0A388JRY3 | DUF2252 domain-containing protein | -1,6415 | 1,6969 | 474 |

**Methods S1 Construction of cultivation box for *Chara braunii***

A commercial thermoelectric cooler (70 L of inner space) was equipped with a custom LED panel (Fig. S1b; composed of RGB, white, UV, plant growth, and aquarium and UV LED types giving individual powers 8.64, 18.00, 4.32, 7.20 and 7.20 W respectively). The spectrum of each LED was measured by STS VIS spectrometer (OceanOptics, slit 25 µm, Fig. S1c). The intensity of each LED type was fully adjustable via Pulse Width Modulation (PWM) control, managed by a Raspberry Pi microcontroller running a Python script available on https://github.com/vosolsob/Cultivation_box. The PWM signal was generated using the pigpio Python module (https://pypi.org/project/pigpio/), which enables hardware timing on the Raspberry Pi's GPIO output interface, resulting in a smoother PWM signal (eliminating LED flicker) compared to a simple software-based approach. The LEDs were powered by a 12V output from a standard ATX PC power supply via N-channel MOSFETs mounted on a custom-made PCB. An internal 80W Peltier element (TEC1-127080S) was powered by the same supply through a Raspberry Pi-controlled N-channel MOSFET and a two-channel relay configured as an H-bridge, allowing for the repolarization of the Peltier element and enabling both heating and cooling modes. The temperature inside the cooler was monitored by two DHT22 sensors placed in opposite corners, with data fed into the Raspberry Pi. The control script operates in an infinite loop, checking the internal temperature every five seconds. The illumination intensity is regulated according to initial parameters such as 'sunrise' and 'sunset' times and the maximum intensity of each LED type. Illumination can be set to remain constant throughout the day or adjusted according to a sinusoidal curve with a 24-hour period. Temperature regulation can either be constant or variable, depending on a three-point setup ('sunrise', 'maximum', and 'sunset' temperatures). At night, the temperature decreases linearly, while during the day, a combination of linear increase between morning and evening points and a sinusoidal component with a half-period equal to daytime is applied. The maximum temperature of this composite curve is adjusted iteratively to match the desired peak temperature, which is typically reached during the cooler's 'afternoon' hours, mimicking natural conditions. The convergence of the actual temperature to the desired value is achieved by gradually adjusting the power of the Peltier element (through MOSFET regulation by PWM) or its repolarization via the H-bridge. While this method of Peltier element regulation using pure PWM without an inductance filter is straightforward to implement, it is not optimal. The element is powered at the maximum voltage during the duty cycle, resulting in greater power losses compared to applying a smoothed voltage. Additionally, if the system is powered by 12V, the maximum power of the element is slightly reduced due to a voltage drop across the

MOSFET. Depending on the temperature regime, the relay in the H-bridge may experience varying degrees of stress, as observed in our system after two years of continuous operation. This issue could be addressed either through software modifications (e.g., brief shutdown of the MOSFET during H-bridge repolarization) or by implementing a semiconductor-based H-bridge. Several levels of security have been implemented in our system. The first level is integrated into the Python script, where an exceedance of 28°C automatically shuts off the LEDs, and a drop below 2°C turns off the Peltier element. The second level of security is at the operating system level: upon startup (or accidental reboot) of the Raspberry Pi, all GPIO ports are automatically set to zero. A monitoring Bash script also runs, checking every minute whether the main Python script is active; if not, it is restarted with parameters loaded from a configuration file. This automation ensures that the control script does not require manual activation when the cooler is powered on. User control of the system is facilitated through a touch display, allowing full access to the Raspberry Pi's operating system.

**Methods S2 Immunolocalization of internodal cells with CbPINs and $H^+$ATPase**

The protocol for immunostaining of internodal cells was modified from (Schmölzer et al. 2011). The thalli of young, elongating *Chara braunii*, strain S276, containing several nodes were cut from its media using scissors and fixed as described in (Schmölzer et al. 2011). Fixed internodal cells of thalli were dissected with a scalpel into fragments 3mm long and further processed as follows: 3x15 min wash with PBS, 30 min treatment with 1 mg $ml^{-1}$ $NaBH_4$ in PBS, 3x15 min wash with PBS. Samples were blocked in 1% (w/v) bovine serum albumin (BSA) and 50 mM glycine in PBS. Primary rabbit antibody against $H^+$-ATPase was used at a dilution 1:1000 (AS07260; Agrisera). Polyclonal antibodies against *Chara* PINa and c, raised in rat against epitopes specific to each PIN protein (Moravian Biotechnology, Brno, Czech Republic) were used in dilution 1:500. Primary antibodies were incubated overnight at 4°C. The next day samples were washed 3x15 min with PBS followed by incubation with secondary antibody, goat anti-Rat IgG (H&L) - Alexa Fluor 488 (A-11006 Invitrogen) for PINs and goat anti-Rabbit IgG (H&L) - Alexa Fluor 546 (A-11035, Invitrogen) for $H^+$-ATPase in 1% (w/v) BSA and 50 mM glycine in PBS for 2h at room temperature. The samples were washed 3x30 min wash with PBS. The last wash was done in sterile water for 10 min, after which samples were placed in 50% glycerol.

**Methods S3 Immunolocalization of antheridia CbPINs and $H^+$ATPase**

Antheridial filaments were immunolocalized using modified protocol from (Żabka et al. 2016). The apical nodes of *Chara braunii* S276 containing generative organs were cut from the rest of the thallus. Cut segments were briefly washed in distilled water that was followed by fixation in 4% paraformaldehyde solution in PBS (pH 7.0) for 45 min with the addition of 0.5 mM $CaCl_2$. Samples were then washed 3x5 with PBS. After that, samples were permeabilized for 10 min in MTSB (50 mM PIPES, 5 mM EGTA, 5 mM $MgSO_4$, pH 7.0; Sigma) containing glycerol (10 %) and Triton X-100 (0.2 %), followed by washing step for 3x5 in MTSB. Samples were treated in ice-cold methanol (-20°C) for 2 min and rehydrated again for 1 min in MTSB. Then cell wall digestion was performed by using 0.1% pectinase from *Aspergillus niger* (Fluka) and 0.01% pectolyase Y-23 (ICN) for 15 min in MTSB which was followed by washing 3x5 in MTSB. Samples were then permeabilized a second time in 10% (v/v) DMSO and 3% (v/v) Nonidet P-40 in MTSB for 1h. After washing 3x5 in MTSB, the samples were placed on superfrost glass (Epredia™ SuperFrost Plus™). The antheridia were dissected from thallus under a binocular microscope, squashed to release the antheridial filaments and air-dried. The slides were then blocked for 1h with 1% BSA in MTSB. The slides were then incubated with a primary antibody containing 1% BSA in MTSB in a humid chamber overnight at 4°C and washed 3x5 with MTSB the next day, followed by incubation with secondary antibody for 2h. The same primary and secondary antibodies were used as in the protocol for immunolocalization of internodal cells (as described above). The samples were washed 3x5 in MTSB. In the last washing step, DAPI was added in a concentration of 1ug $ml^{-1}$. Samples were then washed with sterile water and placed in 50% glycerol.

**Methods S4 Immunolocalization of tubulin in internodal cells**

For the immunostaining of microtubules, thalli were pretreated with perfusion solution (200 mM sucrose, 70 mM KCl, 4.49 mM MgCl2. 5 mM EGTA, 10 mM PIPES, pH=7 (Wasteneys et al. 1989), followed by fixation for 10 min with 1% glutaraldehyde in PBS and cutting off the nodes. After washing in PBS, thalli segments were incubated with 50 mM glycine in PBS and blocked with 1% BSA in PBS for 1h. Primary monoclonal antibodies against α-tubulin (DM1A) (1:1000) in PBS were incubated overnight. The following day, samples were incubated with a secondary, goat anti-mouse IgG, Alexa Fluor™ 488 (A28175), (1:1000) for 3h, after which they were washed with PBS. The final wash was done using a 200 mM sucrose solution.

**Methods S5 Immunolocalization of tubulin in antheridial cells**

Thallus tips containing antheridia were incubated for 1h in a 5 µM taxol solution. Then, the samples were fixed for 40 min in a 1% glutaraldehyde in PMET buffer. After fixation, the samples were washed 3x for 10 min in PMET buffer, followed by cell wall digestion (0.1 %

pectinase from *Aspergillus niger* (Fluka) and 0.01 % pectolyase Y-23 (ICN) for 20 min. After washing the samples were then incubated in a permeabilization buffer for 3h at room temperature and then rinsed three times for 10 min in PBS. Following the washing, samples were incubated with a blocking solution containing 1% BSA in PBS for 60 min. The primary and secondary antibodies step was the same as in internodal cells. Just before the final wash, the samples were stained with 1 µg $ml^{-1}$ Hoechst 33342 for 5 minutes.

**Methods S6 Protein extraction and Western blot**

Around 500 mg of fresh *Chara braunii* thalli strain S276, including rhizoids, was harvested, washed in distilled water, and briefly blotted with a paper towel to remove the excess water. Thalli were homogenized in precooled mortar and pestle with liquid nitrogen. The homogenate was transferred to cold 15 mL centrifuge tube, resuspended in a cold extraction buffer 330 mM saccharose, 100 mM KCl, 1 mM EDTA, 50 mM Tris–HCl pH 7.4, 0.5 mM phenylmethylsulfonyl fluoride (PMSF), 5 mM dithiothreitol (DTT) and 1% (v/v) protease inhibitor cocktail (P9599; Sigma) at a ratio of 0.5 ml $g^{-1}$ FW (according to Schmölzer et al. 2011). Cell debris was removed by centrifugation at 3,000xg for 10 min at 4°C. The pellet was discarded while the supernatant was again centrifuged at 100,000xg for 1h at 4°C. The resulting pellet representing the solubilized membrane fraction was resuspended in the extraction buffer. Protein extracts were mixed in ratio 1:1 with 2D buffer, then separated on a 10% SDS gel electrophoresis. SDS gels were transferred to a nitrocellulose membrane by electro-blotting (Trans-Blot® Turbo™ Transfer System, BioRad). Western blots were probed with rat anti-CbPINa (1:1000), rat anti-CbPINc (1:1000), rabbit anti-AHA (1:2000) and mouse anti-α-tubulin as positive controls, rat pre-immune serum (1:1000) as a negative control, and respective secondary HRP-conjugated antibodies (rabbit anti-rat HRP conjugate ENZO ADI-SAB-200-J 1:5000, goat anti-rabbit HRP conjugate, ENZO ADI-SAB-300-J 1:5,000) Proteins were visualized using the enhanced chemiluminescence (ECL) method (Pierce Western Blotting Substrate) and Azure 600 Imaging System.

**Methods S7 Sample preparation for phosphoproteomic analysis**

Plant material was harvested, frozen in liquid nitrogen, ground to a fine powder, and stored at -80°C until further processing. For comparisons between treatments, all replicates of all treatments were grown on the same day and processed independently.

**Methods S8 Protein extraction for phosphoproteomic analysis**

For protein extraction, samples were suspended in an extraction buffer with 100 mM Tris-HCl pH 8.0, 7 M Urea, 1% Triton-X, 10 mM DTT, 10 U/ml DNase I (Roche), 1 mM $MgCl_2$ and 1% benzonase (Novagen) and lysed by sonication using 30 cycles of 30 seconds ON and 30 seconds OFF at 90% amplitude at 4°C using a waterbath sonicator (Qsonica). Lysate was cleared by centrifugation at 20.000xg for 30 minutes at 4°C. The supernatant was collected and an extra 1% (v:v) of benzonase was added and incubated for 30 minutes at room temperature. Followed by alkylation in 50 mM Acrylamide for another 30 minutes at room temperature. After alkylation, proteins were precipitated using methanol/chloroform. To the one volume lysate, methanol, chloroform, and milliQ were added in a ratio 4:1:3 with rigorous vortexing between each addition. Lysate was centrifuged for 10 minutes at 5000 rpm. After centrifugation, the top layer was discarded and 3 volumes of methanol were added to precipitate the protein layer by centrifugation for 10 minutes at 5000 rpm. After centrifugation, the supernatant was discarded and the protein pellet was air-dried. Next, protein pellets were resuspended in 50 mM ammonium bicarbonate (ABC) and protein concentration was measured by Bradford reagent (Biorad). For every replicate 500 µg protein was digested overnight at room temperature with sequencing grade trypsin (Roche) in a ratio of 1:100 trypsin:protein. After digestion, peptides were desalted and concentrated using homemade C18 microcolumns. Microcolumns were produced using disposable 1000 µl pipette tips that were fitted with 4 plugs of C18 octadecyl 47 mm Disks 2215 (Empore™) material and 1 mg:10 µg of LiChroprep® RP-18 (Merck): peptides. Microcolumns were sequentially washed with 100% methanol, 80% Acetonitrile (CAN) in 0.1% formic acid and twice equilibrated with 5% Acetonitrile in 0.1% formic acid. These steps were performed by centrifugation for 2 minutes at 1500xg. Peptides were loaded onto equilibrated columns for 30 minutes at 400xg. Bound peptides were washed with 5% Acetonitrile in 0.1 % formic acid and eluted with 80% ACN in 0.1% formic acid for 2 minutes at 1500xg.

**Methods S9 Phosphopeptide enrichment**

Phospho-peptide enrichment was performed using PureCube Fe-NTA MagBeads magnetic beads (Cube Biotech) following the manufacturer’s instructions. Eluted peptides were acidified using 10% formic acid. Acidified samples were desalted and concentrated using homemade C18 microcolumns. Microcolumns were produced by fitting disposable 200 µl pipette tips with 2 plugs of C18 octadecyl 47 mm Disks 2215 (Empore™) material and 1mg:10 µg of LiChroprep® RP-18 (Merck): peptides. Microcolumns were washed and equilibrated as described above. Peptides were loaded onto equilibrated microcolumns for 30 minutes at 400xg, 5% acetonitrile in 0.1% formic acid, and eluted with 80% ACN in 0.1% formic acid for 2 min at 1500xg. Eluted peptides were subsequently concentrated using a vacuum concentrator for 30-60 minutes at 45°C and resuspended in 15 µl of 0.1% formic acid.

**Methods S10 Mass spectrometry**

Mass spectrometry was performed as described previously (Kuhn et al., 2024). The MaxQuant quantitative proteomics software package (Tyanova et al. 2016a) was used to analyse LC–MS data with all MS/MS spectra using the following settings: peptide and protein FDR $\leq 0.01$; as protein database the proteome of *Chara braunii* (UniProt ID UP000265515) was used; variable modifications Oxidation (M), Acetyl (protein N-term), Deamidation (NQ), pPhospho (STY); fixed modification AcrylAmide (C); maximum missed cleavage was set at 2; match between runs and label-free quantification options were selected.

**Methods S11 Phosphoproteomic data analysis**

Perseus was used for further analysis of the MaxQuant output PhosphoSTY tab (Tyanova et al. 2016b). Data was imported to Perseus and filtered for reverse and potential contaminants. Phosphosite localization probability was filtered using a cut-off $\geq$0.75. Intensity values were log2 transformed and filtered to contain at least 75% valid values in each group in at least one condition. Values were subsequently normalized by median column subtraction and missing values were imputed from a normal distribution using standard settings in Perseus (width: 0.3, down shift: 1.8). FDR permutation-based t-tests were done in pairwise comparisons (IAA versus DMSO, BA versus DMSO and IAA versus BA). Phosphosites passing the cut -off (FDR $\leq$0.05) were further analyzed.  Adobe Illustrator and R, using standard packages, were used for data visualization.

**Methods references:**

Schmölzer PM, Höftberger M, and Foissner I. Plasma membrane domains participate in pH banding of Chara internodal cells. Plant Cell Physiol. 2011:52(8):1274–1288. https://doi.org/10.1093/pcp/pcr074

Tyanova S, Temu T, and Cox J. The MaxQuant computational platform for mass spectrometry-based shotgun proteomics. Nat Protoc. 2016a:11(12):2301–2319. https://doi.org/10.1038/nprot.2016.136

Tyanova S, Temu T, Sinitcyn P, Carlson A, Hein MY, Geiger T, Mann M, and Cox J. The Perseus computational platform for comprehensive analysis of (prote)omics data. Nat Methods. 2016b:13(9):731–740. https://doi.org/10.1038/nmeth.3901

Wasteneys GO, Jablonsky PP, and Williamson RE. Assembly of purified brain tubulin at cortical and endoplasmic sites in perfused internodal cells of the alga Nitella tasmanica. Cell Biology International Reports. 1989:13(6):513–528. https://doi.org/10.1016/0309-1651(89)90098-2

Żabka A, Polit JT, Winnicki K, Paciorek P, Juszczak J, Nowak M, and Maszewski J. PIN2-like proteins may contribute to the regulation of morphogenetic processes during spermatogenesis in Chara vulgaris. Plant Cell Rep. 2016:35(8):1655–1669. https://doi.org/10.1007/s00299-016-1979-x
